# Human pluripotent stem cell model of multiple epiphyseal dysplasia with *MATN3* mutation identifies altered Matrix organisation and upregulation of the cholesterol biosynthesis pathway

**DOI:** 10.1101/2025.09.01.673425

**Authors:** Steven Woods, Nicola Bates, Stuart Cain, Paul EA Humphreys, Fabrizio E Mancini, Brenda Aguero Burgos, Peter Harley, Rayed Ali A. Alqahtani, Witchayapon Kamprom, Aleksandr Mironov, Antony Adamson, Ian J Donaldson, Geert Mortier, Kate Chandler, Anna Nicolaou, Clair Baldock, Jean-Marc Schwartz, Susan J Kimber

**Affiliations:** Division of Cell Matrix Biology & Regenerative Medicine, Faculty of Biology, Medicine and Health, University of Manchester, Manchester, United Kingdom; Division of Pharmacy and Optometry, Faculty of Biology Medicine and Health, University of Manchester, Manchester, United Kingdom; Electron Microscopy Core Facility, Faculty of Biology, Medicine and Health, University of Manchester, Manchester, United Kingdom; Genome Editing Unit, Faculty of Biology, Medicine and Health, University of Manchester, Manchester, United Kingdom; Bioinformatics Core Facility, University of Manchester; Centre for Human Genetics, University Hospitals Leuven and KU Leuven, Leuven, Belgium; Manchester Centre for Genomic Medicine, Manchester University Hospitals NHS Foundation Trust, Manchester, United Kingdom; Lydia Becker Institute of Immunology and Inflammation, Faculty of Biology Medicine and Health, University of Manchester, Manchester, United Kingdom; Division of Evolution, Infection and Genomics, Faculty of Biology, Medicine and Health, University of Manchester, Manchester, United Kingdom

**Keywords:** Multiple epiphyseal dysplasia, human pluripotent stem cells, cartilage development, osteoarthritis

## Abstract

Multiple epiphyseal dysplasia (MED), caused by mutations in MATN3, is a chondrodysplasia affecting the cartilage growth plate and is characterised by delayed epiphyseal ossification, short stature, and early onset osteoarthritis. Here we generated an in vitro human pluripotent stem cell (hPSC) model of cartilage growth-plate development to identify pathogenic mechanisms underlying MED.

hPSCs were differentiated to chondrocytes via a mesenchymal intermediate, followed by TGFβ3+BMP2 induced chondrogenic pellet culture. MATN3-mutant hPSCs were generated by reprogramming MED patient PBMCs or by CRISPR-Cas9 gene editing to introduce a MATN3 mutation in a hESC line. RNAseq was used to assess chondrogenesis and identify MED pathogenic mechanisms. Transmission electron microscopy (TEM) was used to assess extracellular matrix assembly.

The resultant hPSC-derived cartilage pellets displayed a typical cartilage morphology and strongly expressed cartilage matrix markers, e.g., collagen II and matrilin-3. Matrilin-3 protein was detected within both the matrix and cells of heterozygous mutant hPSC-cartilage pellets. RNAseq of mutant hPSC-cartilage pellets revealed significant enrichment for ‘ECM organisation’ and ‘cholesterol biosynthesis’ pathway genes as well as sightly increased expression of some unfolded protein response (UPR) marker genes. MATN3 mutant hPSC-derived cartilage pellets displayed abnormal matrix assembly, distended ER, accumulation of lipid droplets, and increased cholesterol content.

Our model revealed mutant matrilin-3 induces cholesterol biosynthesis pathway upregulation and abnormal matrix assembly during MED pathogenesis. This study provides new insights into the molecular mechanisms underlying MED and highlights potential therapeutic targets.

## Introduction

Chondrodysplasias are caused by genetic mutations in cartilage growth plate matrix molecules. Due to difficulties in obtaining samples from the developing human growth plate, there is a pressing need for developmental models of conditions affecting the growth plate.

Multiple epiphyseal dysplasia (MED; OMIN 607078) is a chondrodysplasia that causes delayed epiphyseal maturation and early onset osteoarthritis. Patients often require multiple joint replacement surgeries early in life and analgesics to control pain, and there is no cure. There are large variations in phenotype among patients, likely contributed by the specific disease-causing variant each patient has, as well as background genetics. Matrilin-3 is a tetrameric protein containing a von Willebrand Factor A-like domain (vWFA) and four EGF-like domains, which is highly expressed in skeletal tissues especially cartilage (1). In mice it is strongly expressed throughout the growth plate, although more weakly in articular cartilage (2). Monomers have a molecular mass of 53 kDa and form homotetramers, or sometimes heterotetramers with matrilin-1 (3, 4). Mutations in *MATN3*, the gene encoding the matrix protein Matrilin-3 are a cause of MED. with the majority clustered in *MATN3* exon 2 encoding the vWFA domain (5). Matrilin-3 is found in the matrix of cartilage, including prehypertrophic growth plate cartilage during development. It plays a role in crosslinking type II and type IX collagen and other matrix molecules and is therefore proposed to aid the generation of matrix protein networks (6, 7). Moreover, it has been reported to interact with bone morphogenetic protein 2 (BMP2) (8) and increased Matrilin-3 has also been associated with osteoarthritis (9).

Current understanding of MED disease pathogenesis largely comes from mouse models. For example, homozygous knock-in matn3 V194D mutant mice display ER stress/UPR dysregulated apoptosis and reduced proliferation of growth plate chondrocytes (10). ER stress/UPR was also observed following overexpression of mutant matrilin-3 in cell lines (11). Interestingly, humans display an autosomal dominant inheritance (5), whereas mice with heterozygous mutations have a much milder phenotype. There is therefore a need for human MED chondrogenesis models involving endogenous expression of both mutant and wild type (WT) Matrilin-3.

Human pluripotent stem cells (hPSCs) are ideal for generating disease models, due to their virtually unlimited self-renewal and ability to differentiate into multiple cell types, including chondrocytes. Much of the current chondrocyte differentiation research focuses on generating articular-like rather than growth plate-like chondrocytes. Since growth plate cartilage is responsible for elongation of the long bones affected in chondrodysplasias like MED, development of robust growth plate-like chondrocyte differentiation protocols are required for chondrodysplasia disease models.

We developed a pluripotent stem cell to growth plate-like chondrocyte differentiation protocol, resulting in three-dimensional cartilage pellets that display typical cartilage characteristics and closely resemble human fetal limb cartilage development (12). Using both patient derived iPSCs and CRISPR-Cas9 gene editing of one of our previously documented (13-15) human embryonic stem cell (hESC) lines, we implemented our protocol to model MED caused by *MATN3* mutations and uncovered novel pathogenic mechanisms involving activation of the cholesterol biosynthesis pathway and abnormal matrix protein assembly.

## Results

### Differentiation of hPSCs to hypertrophic chondrocytes

To model MED *in vitro* using hPSCs, we first established a robust hPSC to hypertrophic chondrocyte differentiation protocol (Figure 1). Using the hESC line MAN13, we initially differentiated pluripotent stem cells to a mesenchymal intermediate (iMSC) as previously described (16). The iMSCs were generated by switching hPSCs from TeSR-E8 MesenPRO medium, with culture substrate sequentially changed from VTN to gelatine, and then to tissue culture plastic (Figure 1A). Once established, iMSCs could be expanded for around 10 passages while still retaining differentiation capacity. The iMSCs displayed a fibroblast-like morphology and expressed bone marrow mesenchymal stromal cell (BM-MSC) markers CD44, CD73, CD90 and CD105 (Figure 1B and C). Using differentiation medium established for bone marrow (BM)-MSC osteogenesis (OsteoMAX-XF), MAN13-iMSCs underwent osteogenic differentiation in monolayer culture, forming a mineralised matrix by day 28 (Alizarin Red staining) and expressing osteogenic markers RUNX2, ALPL (Alkaline Phosphatase), BGLAP (Osteocalcin), SP7 (Osterix) and SPP7 (Osteopontin) (Figure S1). Interestingly, unlike BM-MSCs, iMSCs failed to undergo adipogenic differentiation using conventional protocols (Figure S2), highlighting key differences from adult BM-MSCs, despite some shared features.

**Figure 1.**
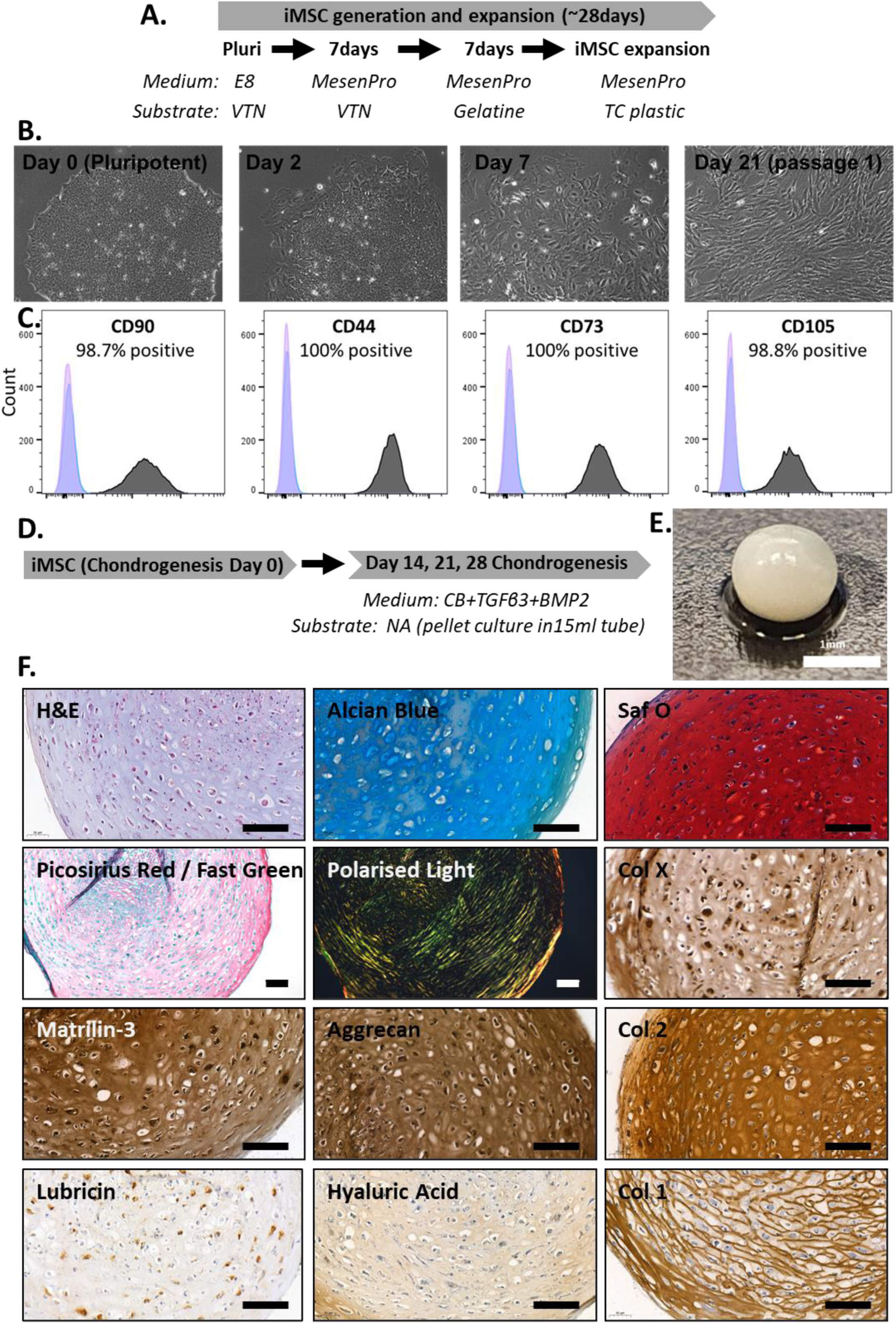
Chondrogenic differentiation of unaffected hESCs. A, Schematic showing protocol for differentiation of hPSCs to hypertrophic chondrocytes via an iMSC intermediate. B, Phase contrast images showing morphological changes during hPSC differentiation to iMSC. C, Flow cytometry analysis of iMSCs for BM-MSC surface markers. Pink represents non-stained (no antibody), blue represents isotype control antibody and grey represents antibody to marker indicated above. D, schematic showing iMSC differentiation to cartilage pellets. E, Photograph of D21 cartilage pellet. F, Histological analysis, immunohistochemistry for matrix components and polarised light microscopy (Picrosirius red stained) for D21 cartilage pellets. Images are representative of 3-5 runs for each of the markers show. Scale bars = 100μm.

Chondrogenic differentiation of iMSCs derived from MAN13 was performed by culturing as cell pellets with addition of 100ng/ml BMP2 and 10ng/ml TGFβ3. Histological analysis of D21 pellets confirmed expression of cartilage-related proteins collagen II, collagen X and matrilin-3, and presence of GAGs (Alcian blue/Safranin O +ve) within an organised collagen matrix (indicated by Picrosirius red and polarised light microscopy) (Figure 1F). RNAseq was performed at D14, D21 and D28 of cartilage pellet culture and compared with iMSCs and the originating pluripotent cells (Figure 2 and S3). Principle component analysis (PCA) showed clear separation of all three stages (Figure 2A). Differential gene expression analysis revealed 7,054 differentially expressed genes (DEGs) between pluripotent and iMSC stage (Figure 2B), significantly enriched for terms such as ‘mesenchyme development’ and ‘ECM organisation’ (Figure 2D). The changes in gene expression are consistent with a loss of pluripotency (e.g. decreased *POU5F1, NANOG, SOX2*) and a gain of mesenchymal (e.g. increased *CD44, NT5E, ENG and ALCAM*) phenotype (Figure 2B). Comparing iMSCs with chondrogenic pellets at days 14, 21 and 28 identified 3,308, 3,767 and 3,797 DEGs, respectively (Figure S3). In addition, direct comparison between pellet stages showed only 9 DEGs between D21 and D14, and 1 DEG between D21 and D28 (Figure S3), illustrating close similarity of D21 transcriptome with both earlier and later pellet stages. We therefore focused subsequent analysis on D21 cartilage pellets. Analysis of the 3,767 DEGs between pellet D21 and iMSC revealed enrichment for terms associated with chondrogenesis (Figure 2C and E). Transcripts significantly upregulated from iMSC stage to D21 cartilage pellet included *SOX9, COL2A1* and *ACAN* which are associated with chondrogenesis in addition to *PTH1R, IHH* and *COL10A1* which are associated with pre-hypertrophic/hypertrophic differentiation and the MED causal gene *MATN3* (Figure 2C).

**Figure 2.**
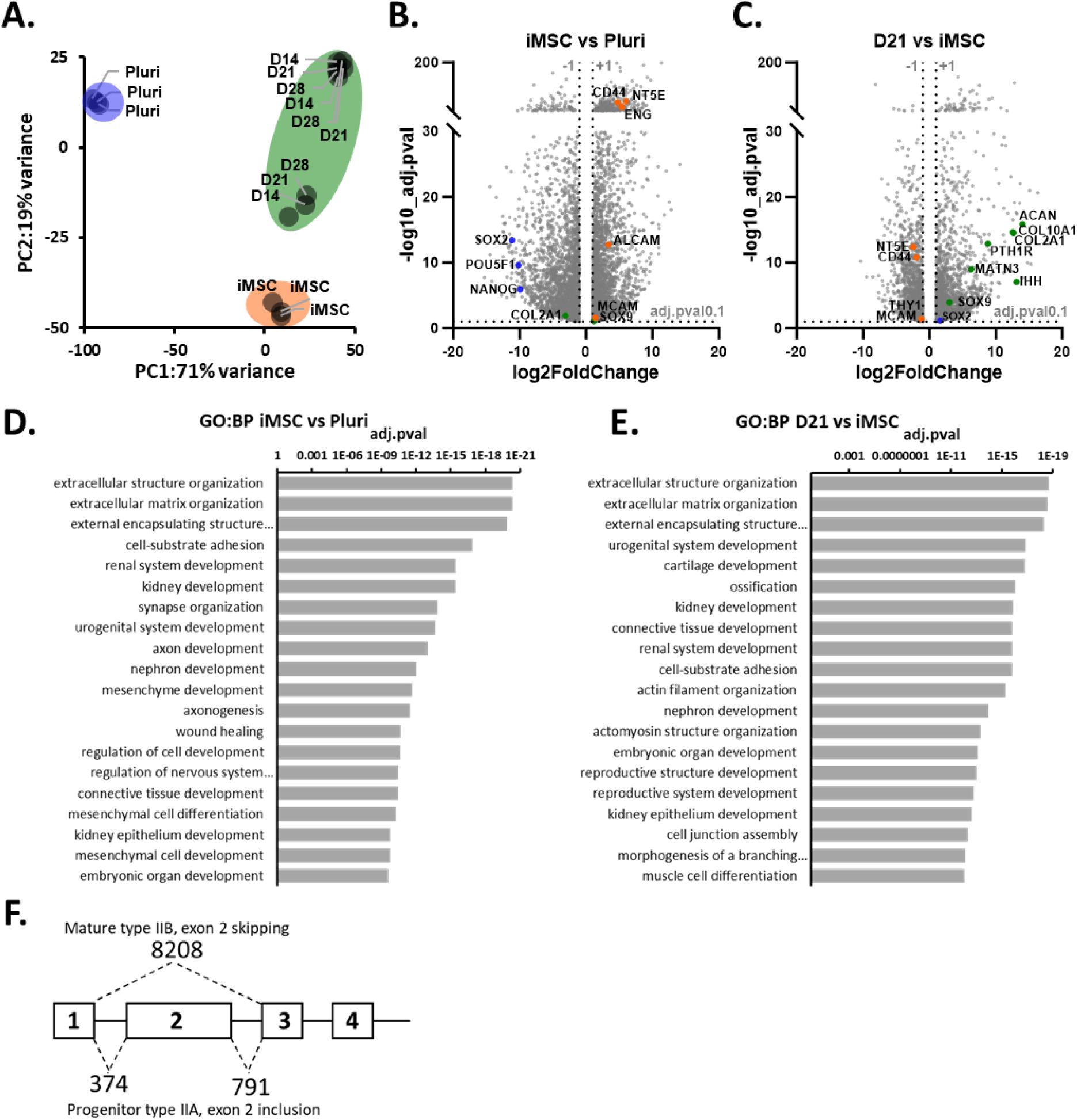
Bioinformatic analysis of unaffected hESCs chondrogenic differentiation. A, principal component analysis (PCA) of cell-transcriptome from different stages and days in the chondrogenic differentiation protocol. B, Volcano plot for genes differentially expressed between iMSC and Pluripotent (pluri) stage of the chondrogenic differentiation protocol. Example DEGs associated with; pluripotency are highlighted in blue, mesenchymal cells are highlighted in orange and chondrogenesis are highlighted in green. C, Volcano plot for genes differentially expressed between D21 and iMSC stage of the chondrogenic differentiation protocol. D, Gene Ontology biological process enrichment analysis for genes differentially expressed between iMSC and pluripotent stage of the chondrogenic differentiation protocol. E, Reaction enrichment analysis for genes differentially expressed between D21 and iMSC stage of the chondrogenic differentiation protocol. F, Alternative splicing analysis for COL2A1 showing mature splice variants. Numbers indicate read count for indicated exon-exon junction. Data combined from 3 independent differentiation runs of the hESC lines MAN13.

Transcript variant analysis of *COL2A1* showed expression of the mature form lacking exon 2 (17), indicating the presence of mature chondrocytes in the D21 cartilage pellets (Figure 2F).

### MED patient iPSC model of chondrogenesis

Peripheral blood mononuclear cells (PBMCs) from four adult MED-patients (two males and one female with V194D and one male with T195K mutation within MATN3) and four unaffected individuals (two males and two female, one being the mother of two V194D patients), were reprogrammed to hiPSCs via erythroid progenitor followed by CytoTune-iPS Sendai based reprograming (18) (Figure S4). Patients had a normal to mildly short stature, experienced joint pain and displayed delayed ossification of the proximal femoral epiphyses (19). Generated HiPSCs were positive for pluripotency-associated markers (Figure S5) and could differentiate to mesoderm, endoderm and ectoderm in embryoid bodies (Figure S6).

All lines formed iMSCs, with minor mutation-independent differences in morphology and CD44, CD73, CD90 and CD105 expression (Figure S7). RNA-seq confirmed loss of pluripotency and gain of mesenchymal identity in both unaffected and affected iMSCs, with most DEGs shared between them (Figure S7C-E). The iMSCs were then differentiated using chondrogenic pellet culture (Figure S8).

RNAseq comparison of the resultant D21 cartilage pellets with their originating iMSCs confirmed increased expression of cartilage-related genes in both unaffected and affected pellets, confirming both lines were capable of chondrogenesis (Figure S8-D). Interestingly, some cartilage genes were increased to a greater extent in affected than unaffected differentiation. Consistent with hESC differentiation (Figure 2A), we observed clear separation by PCA of hiPSC pluripotent, iMSC and D21 cartilage pellets stages (Figure 3B).

**Figure 3.**
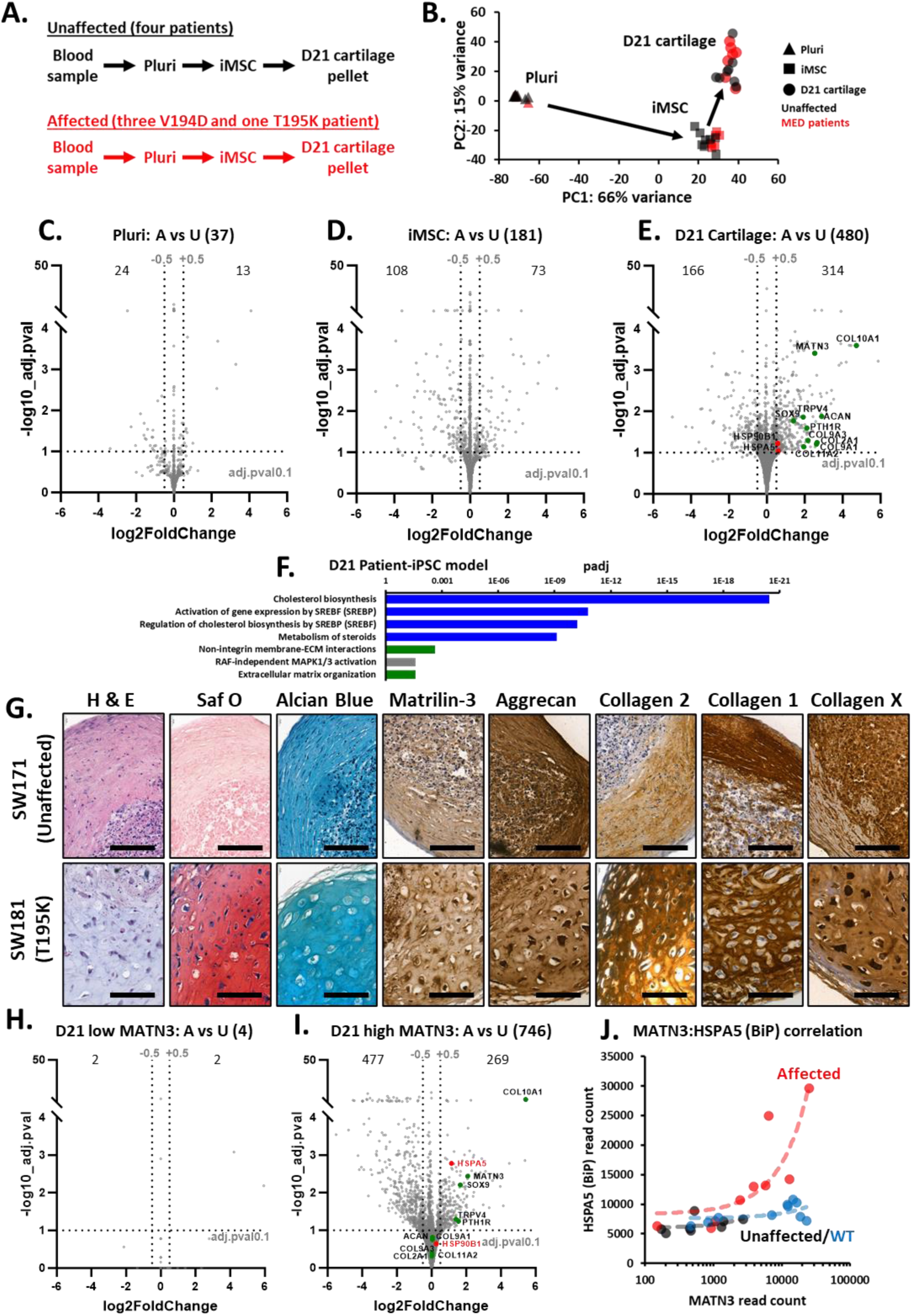
Comparison of affected MED patient with unaffected hiPSC chondrogenesis. A, schematic illustrating experimental design and differentiation of unaffected and MED patient chondrogenesis. B, PCA of RNAseq for unaffected and MED patient chondrogenesis at pluripotent, iMSC and D21 cartilage pellet stages of the chondrogenic differentiation showing segregation by stage. C-E, Volcano plots showing genes differentially expressed when affected MED patient cells (V194D and T195K combined) are directly compared with unaffected cells at each stage of the chondrogenic differentiation protocol. F, Reaction Enrichment analysis for genes differentially expressed between affected MED patient and unaffected D21 cartilage pellets. G, Immunohistological analysis of unaffected and affected MED patient D21 cartilage pellets. Scale bars = 100μm. Note the differences in chondrocyte size. H, Volcano plot showing genes differentially expressed between affected MED patient and unaffected D21 cartilage pellets that express MATN3 transcript below a normalised read count of 1200 (two V194D and one T195K sample vs three unaffected samples). I, Volcano plot showing genes differentially expressed between affected MED patient and unaffected D21 cartilage pellets that express MATN3 transcript above a normalised read count of 1200 (two V194D and two T195K sample vs five unaffected samples). J, Correlation between HSPA5 and MATN3 gene expression in Man13 WT hESC (blue, Pearson correlation coefficient: 0.43), unaffected (black, Pearson correlation coefficient: 0.42) iPSC and affected (red, Pearson correlation coefficient: 0.96) iPSC differentiation.

To investigate MED pathogenesis, we directly compared the transcriptomes of MED patient lines harbouring either V194D or T195K MATN3 (Willebrand factor A domain) mutations with unaffected lines at each differentiation stage. As *MATN3* expression is low at the pluripotent stage, mutations are expected to have minimal impact, however 37 DEGs were still detected, including some genes encoding collagens and other matrix proteins (Figure 3C). At the iMSC stage we identified 181 DEGs between affected and unaffected cells (Figure 3D). *MATN3* expression is highest at the cartilage pellet stage (Figure 2), therefore, as expected, this was the stage at which we observed the greatest number of DEGs (480) between affected and unaffected cells (Figure 3E), indicating this is the stage most impacted by MATN3 mutations. Analysis revealed mutant enrichment of genes associated with ‘Cholesterol biosynthesis’, ‘Extracellular matrix organisation’ (Figure 3F) and chondrogenesis e.g. *COL2A1, SOX9, COL10A1* and *MATN3* itself (Figure 3E). Consistent with previous models of MED we identified a significant (padj<0.1), although modest, increase in the protein chaperone and UPR markers HSPA5 (BiP) and HSP90B1 (GRP94) (Figure 3E). Pellet size differed by donor rather than by presence or absence of mutation. We observed a range of chondrocyte morphologies with a tendency for those in affected pellets to be larger (Figure 3G and S8).

To test if the mutant phenotype depends on MATN3 levels, pellets were grouped by high or low MATN3 transcript expression, and differential expression analysis was performed between affected and unaffected samples within each group. Only 4 DEGs were found in the low *MATN3* expression group whereas 746 DEGs were found in the high MATN3 expression group (Figure 3H and I), confirming high expression of mutant *MATN3* is responsible for driving differences between affected and unaffected cartilage pellets. Examining the correlation between *HSPA5* and *MATN3* expression (Figure 3J), we found that the UPR marker *HSPA5* was increased to a greater extent in high *MATN3* expressing pellets (Figure 3I-log2FC 1.14) than when all pellets were included in the analysis (Figure 3E-log2FC 0.60).

### Using CRISPR-Cas9 genome editing to generate an hESC model of MED

To validate findings from the MED patient derived iPSC model, we generated isogenic hESC lines differing only by the heterozygous MATN3 V194D mutation, which is present in three of four patients and also previously modelled in mice (20) (Figures 1 and 2). To create the mutation, we used a CRISPR-Cas9 homology directed repair (HDR) method that included a targeting repair vector harbouring the *MATN3* V194D (rs104893645 A>T) mutation and loxP-CRE removable pGK driven mCherry selection flanked by 5’ and 3’ homology arms (Figure 4A and S9). Following nucleofection, cells were sorted twice for mCherry positive cells, firstly to enrich for cells that had taken up the plasmid and secondly for cells that had stable expression of mCherry. Following validation of correct targeting vector integration into *MATN3* (Figure S9), cells were nucleofected and sorted for CRE-GFP (Figure S9). After 2 subsequent passages cells were negative for both GFP and mCherry indicating successful removal of the selection marker (Figure S9). Following clonal selection, the presence of a heterozygous mutation was validated using Sanger sequencing (Figure 4A).

**Figure 4.**
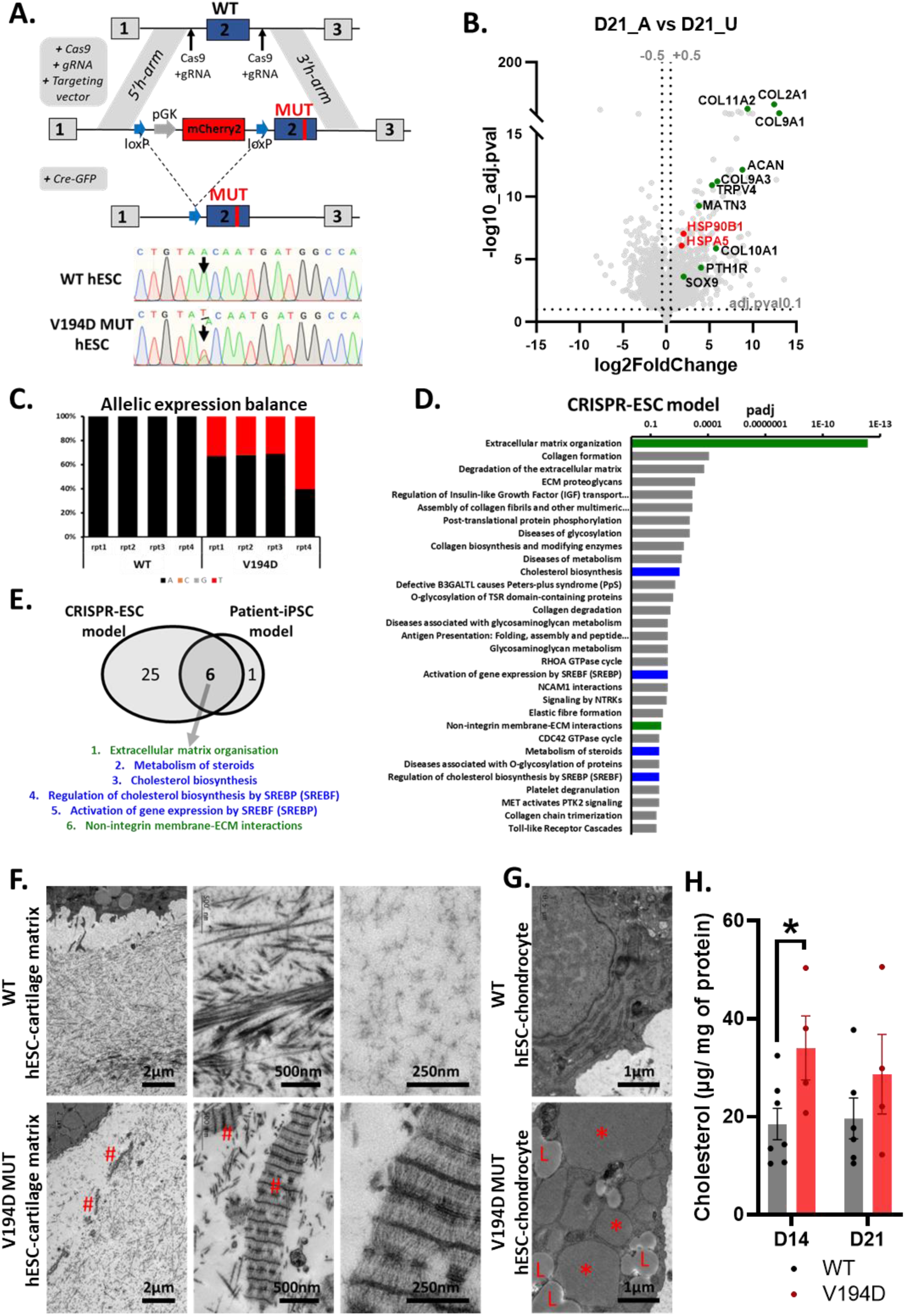
CRISPR-Cas9 hESC model of MED recapitulates patient iPSC model. A, strategy used to create a heterozygous MATN3 V194D mutation in the MAN13 hESC line. B, Volcano plot comparing RNAseq from V194D mutant CRISPR-Cas9 hESC derived cartilage pellets with WT cartilage pellets. C, Transcript expression balance between WT and V194D mutant alleles of WT and V194D mutant pellets. D, Reaction enrichment analysis for genes differentially expressed between WT and V194D (CRISPR-Cas9 mutant) D21 cartilage pellets. E, Venn diagram to identify Reactome terms enriched in both affected MED-patient iPSC-derived D21 cartilage pellets and CRISPR-Cas9 mutant hESC-derived D21 cartilage pellets. F. Transmission electron microscopy (TEM) of D21 cartilage pellet matrix. In the V194D mutant, ‘#’ marks altered matrix structures in the pericellular region at low magnification (left and middle panels). The right panel shows a zoomed-in view of these altered structures. The equivalent pericellular region in the WT is shown for comparison. G, TEM of chondrocytes with D21 cartilage pellets ‘*’ indicates presence of distended ER, ‘L’ indicates lipid droplet accumulation. Electron microscopy was performed at day 21 on two V194D mutation (CRISPR isogenic) and two WT differentiation runs. Data shown are representative of >20 cells from each differentiation runs. H, Quantification of cholesterol in V194D mutant CRISPR-Cas9 hESC derived cartilage pellets compared to WT cartilage pellets using gas chromatography, data combined from three V194D and three WT differentiation runs, statistical differences were calculated using independent two-sample t-test.

Heterozygous V194D mutant MAN13 hESCs were differentiated alongside their isogenic control using our chondrogenesis protocol. MATN3 was expressed approximately equally from the WT and mutant alleles of the CRISPR edited cells, further confirming successful generation of the V194D mutation (Figure 4C). RNAseq of D21 cartilage pellets revealed 1978 DEGs between V194D mutants and WT controls (Figure 4B). Consistent with the findings from patient derived iPSC lines (Figure 3) we again identified increased expression of genes associated with chondrogenesis e.g. COL2A1, SOX9, COL10A1 and MATN3, as well as modest increases in the UPR markers HSPA5 (BiP) and HSP90B1 (GRP94) (Figure 4B). Enrichment analysis replicated the findings in our patient-iPSC model, showing significant enrichment for ‘Cholesterol biosynthesis’, ‘Extracellular matrix organisation’ and related terms (Figure 4D and E and S13 A &B).

Having established that changes in matrix transcripts were consistent across patient-iPSC and CRISPR-hESC models of MED, we investigated if there were changes in matrix protein organisation. Histological analysis indicated cellular retention of matrilin-3 in mutant pellets (Figure 4F S10), as in patient cartilage (Figure 3G). Transmission electron microscopy (TEM) revealed substantial differences in matrix organisation. The pericellular region of V194D mutant pellets were mostly devoid of a diffuse grainy material found in unaffected pellets. Instead, highly organised banded structures reminiscent of type VI collagen were prevalent (Figure 4F and S11). This change in matrix organisation was in keeping with our transcriptome enrichment analysis of hESC and patient iPSC mutant pellets (Figure 4D and 3F) indicating changes in matrix organisation. TEM also revealed distended ER (Figure 4G and S11) consistent with upregulation of ER chaperone gene *HSPA5* (Figure 3E, 3I and 4B). Furthermore, affected day 21 chondrocytes contained many lipid droplets, not seen in unaffected chondrocytes (Figure 4G). Given the observation that cholesterol biosynthesis genes were upregulated at a transcript level, we measured the amount of cholesterol in pellets using gas chromatography. Indeed, mutant pellets contained significantly more cholesterol than WT at day 14 (Figure 4H).

Regulation of expression of cholesterol biosynthesis genes is tightly controlled through multiple factors. To confirm that activation of this pathway is a direct consequence of mutant matrilin-3 protein expression we developed a TC28a2 cell line carrying dox-inducible mutant matrilin-3 (Figure S12; TC28a2-dox-MATN3-V194D cells). Dox treatment led to strong induction of cellular mutant matrilin-3 (Figure S12B), followed by increased expression (and cleavage) of the critical transcription factor SREBP2, and protein expression of its response gene the low-density lipoprotein receptor (LDLR) (Figure S12C-D).

### Bioinformatic comparison of patient iPSC-, CRISPR hESC- and mouse-models of MED

There are few published models of MED, but we wanted to determine if these also involved changes in matrix organisation and activation of the cholesterol biosynthesis pathway. By reanalysing microarray data for the previously published V194D homozygous mouse model, we identified a strong enrichment of the proteins involved in the UPR (Figure S13D), consistent with the original analysis (20). In addition, we also identified a significant enrichment for cholesterol biosynthesis (Figure S13D). Similarly, re-analysis of microarray from a *MATN3* T120M mutant human iPSC model (21) also revealed enrichment for UPR and cholesterol biosynthesis (Figure S13C). Reactome enrichment analysis across the four studies identified five shared terms, one for matrix organisation and four for cholesterol biosynthesis (Figure 5A). UPR/ER stress related terms were only enriched within the homozygous mouse and the T120M iPSC models (21), but did not reach significance in our patient iPSC or CRISPR-hESC models. Notably, most genes associated with cholesterol biosynthesis and UPR show increased expression in all four models, although some changes are modest (Figure 5C, S13A-D and S14).

**Figure 5.**
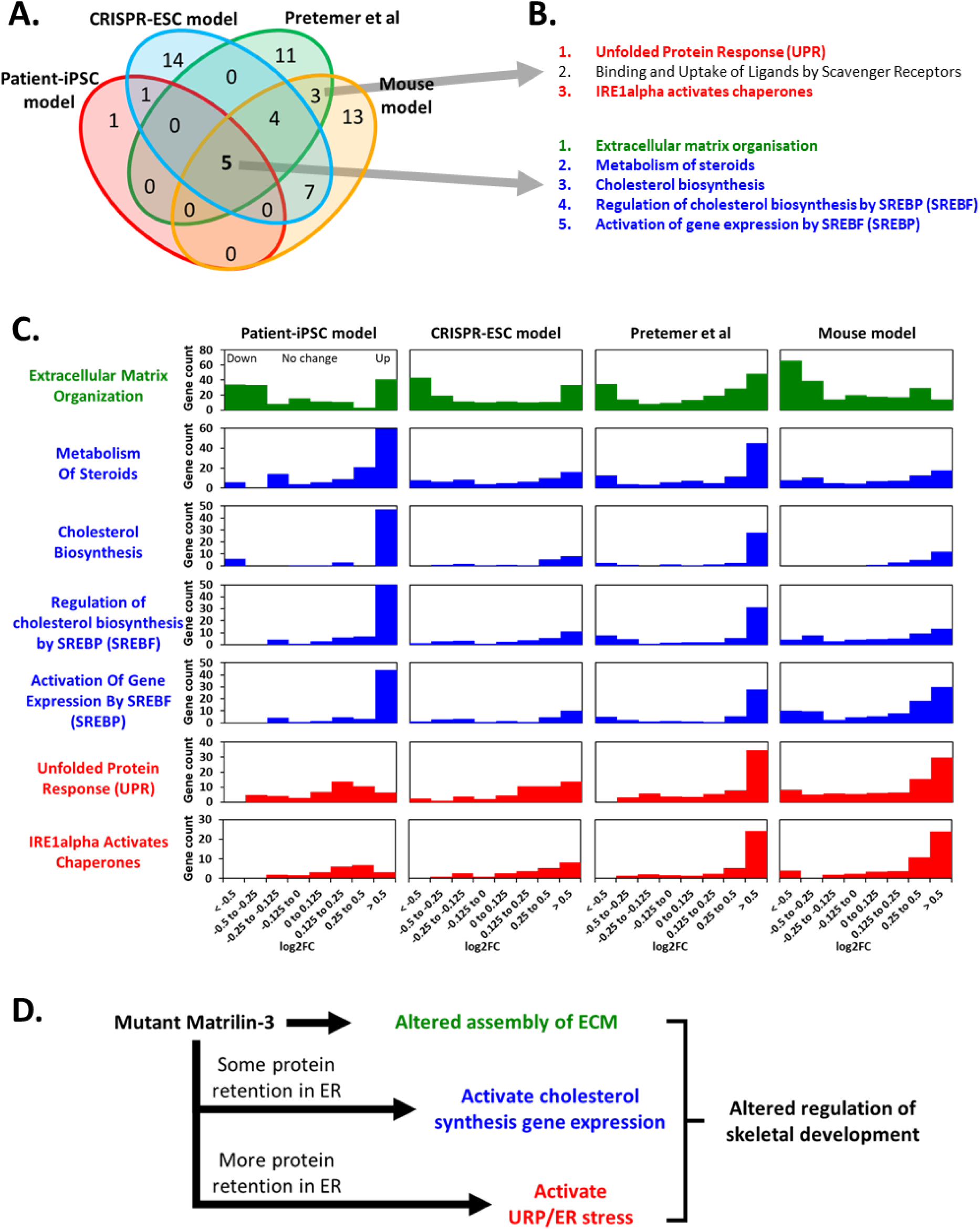
Comparison of patient iPSC model and CRISPR hESC model with previously published models of MED. A, Venn diagram of Reactome pathways enriched within affected/mutant cartilage from either patient iPSC model (Figure 2), CRISPR hESC model (Figure 4), heterozygous MATN3 T120M mutant human iPSC model (21) or homozygous Matn3 V194D mutant mouse model (20). B, list of Reactome pathways that are common to all four models or found only in botj (21) and (20). C, Histogram showing Reactome pathway gene count per 1000 genes for genes expression of which changed by indicated log2FC. Note the varying level of enrichment for both cholesterol and UPR related genes within upregulated genes of each of the four models. D, Schematic illustrating the proposed model by which mutant matrilin-3 alters chondrogenesis and skeletal development in MED patients

## Discussion

The role of matrilin-3 in human cartilage development and disease is unclear, with conflicting findings from earlier studies. For example, Ko et al showed no skeletal abnormalities in matrilin-3 deficient R1/C57Bl/6 mice (22), while van der Weyden et al. showed that knock-out on a 129/C57 background caused premature hypertrophy and cartilage degradation, despite no obvious evidence of chondrodysplasia (23). Leighton et al found slight shortening of the long bones and growth plate dysregulation in homozygous *matn3* mutant mice, which replicates human MED more faithfully than the full knockouts (10). Notable, all these models involved disruption of both *Matn3* alleles, in contrast to human MED caused by heterozygous *MATN3* mutations.

Here we used our protocol for human pluripotent stem cell differentiation to hypertrophic-like cartilage through a mesenchymal intermediate (19), to investigate the chondrogenic phenotype in MED caused by heterozygous mutations within the β-sheet region of the matrilin-3 von Willebrand A-domain. The three-dimensional cartilage pellets produced, stained strongly for type 2 collagen, aggrecan and matrilin-3 indicating a strong cartilage phenotype. Some areas within the pellets also expressed type 1 collagen, lubricin and type 10 collagen indicating chondrocytes at different stages of development. The pellets expressed transcripts characteristic of chondrocytes e.g. *SOX9*, *COL2A1*, *ACAN* and *MATN3* as well as growth plate chondrocyte-specific markers including *PTH1R*, *IHH* and *COL10A1* by day 21 of the protocol. Although these iMSCs share some BM-MSC surface markers and osteogenic potential, they differ in their inability to undergo adipogenesis.

To avoid issues caused by donor age or genetic background we examined iPSC-chondrogenesis in four patient iPSC lines (three with V194D mutations, one with T195K mutation) and an isogenic CRISPRed V194D mutant line. Finding were also compared to previously published MED models to identify common pathogenic mechanisms.

As matrilin-3 is a cartilage matrix protein, it is not surprising that we identified gene expression differences in matrix organisation including upregulation of genes associated with chondrogenesis e.g. *SOX9*, *COL2A1*, *ACAN*, *COL9A1, COL11A2, TRPV4*, *PTH1R*, *COL10A1* and *MATN3* itself in our transcriptome analysis of both affected patient-iPSC and mutant CRISPR-hESC chondrogenesis. This may indicate feedback regulation.

Matrilin-3 binds indirectly to collagen IX in murine cartilage (24) and to collagen VI via the proteoglycans biglycan and decorin (25), so alterations in its tetramer structure and function are likely to disrupt the extracellular matrix ultrastructure. Here we show matrilin-3 mutant hPSC-chondrocytes form abnormal banded structures in cartilage, closely resembling disordered collagen architecture seen in pathological tissues. Often, these are attributed to pathological assemblies of collagen VI, termed Luse bodies or zebra collagen (26) and observed in Bruch’s membrane of the eye in age-related macular degeneration (AMD)(27). Loss of functional matrilin-3 may allow spurious collagen VI assembly into microfibrils. However, unlike the double beaded arrangement of collagen VI microfibrils with typical 105nm periodic dark and light banding, our structures showed 235 nm spacing. This may be due to the binding of additional molecules in these arrays, as seen in AMD patient Bruch’s membrane. There, abnormal collagen VI banding, lacking regularly spaced densities, was linked to such interactions (27). Interestingly, protein analysis of V194D mutant mouse cartilage suggested differences in the associations of other matrix proteins in cartilage (28). It has been shown that the MATN3 V194D mutation causes matrilin-3 to form high molecular weight aggregates (11).

Other collagens have also been linked to formation of zebra, or fibrous long-spacing (FLS) collagen, with variable periodicities (29) e.g. formation of FLS collagen with longer periodicity from assemblies of collagen II, in limb bud cultures upon addition of highly sulphated GAGs (30). FLS collagen was also observed in the skin of a patient with X-linked spondyloepiphyseal dysplasia tarda (31) and in the pleura of a patient with malignant mesothelioma (32). The aggregates are likely caused by imbalance in collagen binding proteins or proteoglycans and identification will require further work. Moreover proteolysis (33) or metal ion binding to the von Willebrand domain, may be affected by the mutation, altering matrilin-3 turnover and matrix cross linking.

We also observed increased MATN3 gene expression in mutant lines, suggesting a feedback response to matrix disruption driving enhanced matrix synthesis. Interestingly, matrilin-3 levels in newborn heterozygous mutant mice appear reduced, however by day 21, staining appears comparable to, or greater than, in wild-type mice (10).

Consistent with previous homozygous mutant *matn3* MED models (10, 20) we observed ER distension along with matrilin-3 and possibly other matrix protein retention (21), but unlike those studies, we saw no marked ER stress or unfolded protein response. Our data suggest these differences relate to mutant matrilin-3 expression, for example, we cultured pellets for 21 days, whereas Pretemer et al used 56 days. Despite protocol differences, we hypothesise that longer culture allowed more mutant matrilin-3 accumulation and UPR activation in their study (21).

Nundlall et al (2010) analysed 5-day-old rib cages from homozygous *matn3* V194D mutant mice expressing only mutant matrilin-3. In contrast, our human cells carry a heterozygous mutation and express both mutant and wild-type protein, as in patients with MED. The presence of wild-type protein, likely forming mixed tetramers with mutant protein, may enhance folding, delaying UPR activation, and reduce the impact compared with fully mutant tetramers. Notably, some MED-causing *COMP* mutations do not cause UPR in mouse models (11), suggesting MED pathogenesis may involve multiple mechanisms.

We observed a significant activation of the cholesterol biosynthesis pathway, confirmed by increased levels of cholesterol in mutant cartilage pellets. Sterol regulatory element-binding proteins (SREBPs) regulate cholesterol pathway gene expression, by responding to lipid levels in the ER (34). When lipid content is normal, INSIG inhibits SCAP (SREBP cleavage-activating protein) mediated SREBP transport to the Golgi. When ER sterols are low SCAP mediates SREBP transport to the Golgi. In the Golgi, Site-1 and Site-2 proteases cleave SREBPs, giving rise to active transcription factors which then translocate to the nucleus and increase cholesterol synthesis gene expression (35). We hypothesize that mutant matrilin-3 disrupts this process. Indeed, overexpression of mutant matrilin-3 in a chondrogenic line led to increased SREBP cleavage and more intense staining for LDLR, a gene previously shown to be transcribed in response to SREBP nuclear signalling (36). SREBF2 (aka SREBP2) and LDLR, along with other genes in the cholesterol synthesis pathway, were also upregulated in *MATN3* mutant hiPSC-derived cartilage pellets. *SREBF2* (log₂FC 0.88, padj 0.005) was significantly increased in both the patient iPSC model (log₂FC 1.49, padj 0.00011) and the CRISPR-edited hESC model, whereas *LDLR* (log₂FC 2.15, padj 0.006) was only significantly upregulated in the patient iPSC model. If confirmed in patients, this dysregulation of steroid synthesis may contribute to symptoms and presents new possibilities for therapeutic intervention and for monitoring disease progression.

Here we used hPSCs to model MED caused by V194D and T195K mutations in matrilin-3, comparing findings to a homozygous V194D mutant mouse model (20) and a hPSC model of the T120M mutation (21). As all these mutations reside within a β-sheet region of the matrilin-3 A-domain (37), it is likely that they induce common pathogenic mechanisms. Thus, we propose a model whereby mutant matrilin-3 will alter matrix organisation and active cholesterol biosynthesis pathway gene expression, then later, or once mutant molecules have accumulated in large enough quantities, it may activate the UPR (Figure 5).

## Methods

### Human embryonic stem culture

HESC line Man13 (15) was cultured as reported previously, see Supplementary Methods. HESCs and hiPSCs were differentiated to iMSCs as previously reported (16) and detailed in Supplementary Methods.

### Osteogenic and Adipogenic differentiation of iMSCs

Osteogenic differentiation of iMSCs was performed using OsteoMax (Millipore). Adipogenic differentiation of iMSCs and BM-MSCs was based on previously published protocols (38) see Supplementary Methods.

### Chondrogenic differentiation of iMSCs

Passage 3 to 5 iMSCs were differentiated to chondrocytes in cartilage pellet culture. iMSCs were dissociated using TrypLE, washed with PBS, counted, and then resuspended in chondrogenic medium at a density of 200k cells/ml. Chondrogenic medium was composed of DMEM, L-glutamine, 100mM dexamethasone (Sigma D4902), 50µg/ml ascorbic acid 2-phosphate (Sigma A8960), 40μug/ml L-proline (sigma P5607), 1x ITS+l (Sigma I2521), 10ng/ml TGFβ3 and 100ng/ml BMP2.

Pellets were then formed by centrifuging 1ml of cell suspension (200k) for 4min at 400g in Corning 15ml centrifuge tubes. Following centrifugation the cap of each tube was loosened by half a turn to allow gas exchange. Cartilage pellets were incubated at 37°C and 5%CO_2_ in air. After 24h pellets were agitated manually to ensure detachment from the tube and then fed every 3 days with fresh chondrogenic medium. At day 21, unless otherwise stated in figure, pellets were then fixed with 4%PFA for histology, fixed with 4% formaldehyde and 2.5% glutaraldehyde in 0.1M HEPES buffer (pH 7.2) for electron microscopy or RNA was harvested for RNAseq.

### Reprogramming of PBMCs to iPSCs

Erythroid progenitors expanded from peripheral blood mononuclear cells (PBMSs) were reprogrammed to iPSCs using CytoTune™-iPS 2.0 Sendai, see Supplementary Methods.

### RNA sequencing and bioinformatic analysis

RNA sequencing and bioinformatic analysis was performed as documented in Supplementary Methods.

### Comparison to existing models of MED

Reactome enrichment analysis from our heterozygous patient hiPSC and heterozygous CRISPR hESC models was performed as described in Supplementary Methods and compared with Reactome enrichment analysis for a homozygous matn*3*V194D mouse model (20) and a heterozygous MATN3 T120M hiPSC model (21). For homozygous matn3V194D mouse model, micro array data previously described (20) was analysed. Probes corresponding to mouse genes (ENSMUSG ID) were converted into human genes (ENSG ID) from Ensembl v103. Probes that did not match to a human ortholog gene were removed, leaving 28,050 probes corresponding to 15,116 human ortholog genes and used as the background for mouse enrichment analysis. Probes with PPLR >0.95/<0.05 were used for Reactome enrichment analysis (1,727probes/1,556 genes). Reactome enrichment analysis was then performed as described in Supplementary. Microarray data for the MANT3 T120M mutant and isogenic control (GSE148728) hiPSC model were downloaded from GEO (NCBI) and analysed using Limma. Adjusted p values were calculated using Benjamini & Hochberg method with the GEO2R program. Probe IDs were filtered for those corresponding to genes with ENSG ID within the Ensembl v103, leaving 20,383 genes in total with 428 DEGs (padj <0.05, log2FC >0.5 or <-0.5). Reactome enrichment analysis was then performed as described above. Lists of significantly enriched pathways (adjusted p value <0.05) for each of the four MED models were then compared to identify pathways in common.

### CRISPR-Cas9 gene editing

CRISPR-Cas9 gene editing to create the MATN3 V194D mutation was performed using a modified exon replacement strategy based on (39). See Supplementary Methods for details.

Immunocytochemistry of cultured cells, embryoid body formation, Flow cytometry, RNA extraction and RT-qPCR, histological analysis and immunocytochemistry of paraffin embedded pellets, Western blotting, electron microscopy, cholesterol analysis and generation of doxycycline inducible mutant Matrilin-3 in TC28a2 cell line are documented in Supplementary Methods.

## Data statement

The data presented in this study are available in the ArrayExpress database at EMBL-EBI.

## Acknowledgements

We thank David Chapman of the UoM (University of Manchester) flow cytometry facility for assistance with cell sorting, Hayley Bennet of the UoM GEU for assistance with cloning MATN3 V194D mutant repair template, Leo Zeef (UoM Bioinformatics Facility) for assistance with data analysis and Jamie Soul (UoM) for assistance with data analysis. We also thank Prof Frank Zauck for providing antibodies to matrilin-3. We are grateful to Prof Mike Briggs of Newcastle University for useful discussions and for providing access to mouse model microarray data.

## Author contributions

**SW** Conception and Design, Data Acquisition, Analysis and Interpretation of Data, Drafting the Article, Critical Revision of the Article, Obtaining of funding. **NB** Data Acquisition, Analysis and Interpretation of Data, Drafting the Article. **PEAH** Data Acquisition, Analysis and Interpretation of Data, Drafting the Article, Critical Revision of the Article. **FEM** Data Acquisition, Analysis and Interpretation of Data, Drafting the Article. **BAB** Data Acquisition, Critical Revision of the Article. **PH** Data Acquisition, Analysis and Interpretation of Data, Drafting the Article. **RAAA** Data Acquisition, Analysis and Interpretation of Data, Drafting the Article. **WK** Data Acquisition, Drafting the Article. **AM** Data Acquisition, Analysis and Interpretation of Data, Drafting the Article. **AA** Drafting the Article, Analysis and Interpretation of Data. **IJD** Analysis and Interpretation of Data, Critical Revision of the Article. **GM** Conception and Design, Provision of patients. **KC** Conception and Design, Provision of patients. **AN** Analysis and Interpretation of Data, Critical Revision of the Article. **CB** Analysis and Interpretation of Data, Drafting the Article, Critical Revision of the Article, Obtaining of funding. **JMS** Analysis and Interpretation of Data, Critical Revision of the Article. **SJK** Conception and Design, Analysis and Interpretation of Data, Drafting the Article, Critical Revision of the Article, Obtaining of funding. **All authors** Final approval of article.

## Funding

This research was funded by the European Union (SYBIL European Community’s Seventh Framework Program) and the UK Medical Research Council (MR/S002553/1 and MR/X002020/1).

## Competing interest statement

The authors declare no conflicts of interest.

## Notes

### Competing Interest Statement

The authors have declared no competing interest.

