## Supplementary methods for "Human pluripotent stem cell model of multiple epiphyseal dysplasia with *MATN3* mutation identifies altered Matrix organisation and upregulation of the cholesterol biosynthesis pathway"

**Human embryonic stem culture**

The human embryonic stem cell (hESC) line 'Man13' (https://hpscreg.eu/cell-line/UMANe002-A) (13) was cultured as colonies under feeder-free conditions in TeSR™-E8™ culture medium (StemCell Tech, Catalogue # 05990), on 6-well tissue culture plates pre-coated with VTN (GIBCO 14700). Stem cell colonies were passaged every 7 days and before reaching ~50-80% confluence using 0.5 mM EDTA (life tech 15575-038).

**Reprogramming of PBMCs to iPSCs**

Peripheral blood mononuclear cells (PBMCs) were obtained from peripheral blood samples donated under local ethical approval and fully informed consent (REC 13/EM/0388 IRAS ID 11469 or REC 11/H1003/3; IRAS ID 64321). Peripheral blood was collected into an EDTA vacutainer. and kept at room temperature until processing within 3 days. To extract PBMCs, blood was mixed 50:50 with PBS w/o calcium and magnesium, carefully layered over Ficoll® Paque Plus and centrifuged at 400g for 40min. PBMCs were then transferred to a new tube and washed twice with PBS. Isolated PBMCs were then frozen in PSC cryomedium at around $1x10^6$ cells per vial or approximately one vial per ml of blood. PBMCs were expanded to erythroid progenitors using StemSpan™ SFEM II + StemSpan™ Erythroid Expansion Supplement, (StemCell technologies). Starting with $0.5x10^6$ PBMCs per well of a 6 well plate, after 24h all non-adherent cells were passaged to a fresh well. After an additional 24h an 80% media change was performed, repeated after an additional 2days and 4 days.  At day 8 post thaw 50k cells were transduced with  CytoTune™-iPS 2.0 Sendai at an MOI of '553' (MOI of 5 for KOS and hc-Myc vectors and MOI of 3 for hKlf4 vector) in 250µl complete erythroid expansion medium and centrifuged at 300g for 35min. Following centrifugation 150µl complete erythroid expansion medium was added giving  400µl , divided as 200µl per well in 2 wells of  a non-coated-standard 24 well TC plate), and incubated for 24h after which 200ul complete erythroid expansion medium was added/well. After an additional 48h a further 200ul was added giving a total of 600µl per well. After a further 24h (day 12 post thaw), cells were transferred to vitronectin (VTN N- terminal truncated (Gibco) coated 6 well plates containing 500µl ReproTeSR (StemCell Technologies) giving a (total 1.1ml /well. After 48h incubation 1ml ReproTeSR was added (total 2.1ml/well. After an additional 24h some cells adhered to the VTN substrate, and all medium was removed and replaced with 1.5ml ReproTeSR. Cells were fed every 24h with 1.5ml ReproTeSR until colony isolation at around day 26-30 post PBMC thaw. Colonies with pluripotent morphology and large enough to handle were isolated using a pulled glass pipette.and maintained in TeSRE8 until a minimum of passage 15 prior to differentiation.

**Human embryonic stem differentiation to iMSCs**

Four days prior to starting differentiation human embryonic stem cell colonies were passaged using EDTA and plated to produce ~10 colonies per well of a VTN coated 6 well plate. To induce differentiation of hESCs to iMSCs, culture medium was switched from TeSR™-E8™ to MesenPRO RS™ medium (Gibco 12746012). MesenPRO RS™ medium was then replaced every 2days. 7days after switching to MesenPRO RS™ medium, cells were passaged using TrypLE (GIBCO 12604-021) and re-plated into a T75 culture flask precoated in a 0.1% gelatine solution (Sigma G1393). After a further 7 days of growth in MesenPRO RS™ medium, cells were passaged using TrypLE and replated into T75 tissue culture flasks (no additional coating) at a split ratio of 1:8, this is referred to as P1 iMSC. iMSCs were then maintained in T75s using MesenPRO RS™ medium and passaged at 1:8 split ratio approximately every 5days or at ~80% confluence using TrypLE.

**Osteogenic differentiation of iMSCs**

Osteogenic differentiation of iMSCs was performed using OsteoMax (Millipore) in 24 well plates. Once cells had reached confluence, medium was switched from MesenPro to

Osteomax. Osteomax medium was then changed every 3 days. Cells were then fixed for Alizarin Red staining or harvested for RNA at days 1, 3, 7, 14 and 28.

**Adipogenic differentiation of iMSCs and BM-MSCs**

Adipogenic differentiation of iMSCs and BM-MSCs (Lonza) was performed using adipogenic medium in 24 wells plates. Adipogenic medium was composed of DMEM, 10%FBS, pen-strep, l-glutamine, 100nM dexamethasone, 10ug/ml IBMX (Sigma I7018) and 10.2ug/ml insulin (Sigma I9278). Medium was then changed every 3 days until Day 21 and 28 when cells were fixed for Oil Red O staining.

**Immunofluorescence**

Pluripotent colonies were passaged onto VTN coated 24well plates using EDTA. Cells were washed twice in PBS and then fixed in 4% paraformaldehyde (PFA) for 20 minutes, followed by another two PBS washes. The fixed cells were blocked and permeabilised for 30min with 3% bovine serum albumin (BSA)/0.3% Triton-X in PBS before overnight incubation at 4˚C with primary antibodies for Nanog, Oct4, SSEA1, SSEA4 and TRA160 diluted in 3% BSA/PBS. They were then washed three times with PBS/0.1% Triton-X, followed by Alexa-Fluor™-488- or Alexa-Fluor™-594-labelled, species-specific secondary antibodies (Life Technologies; 1:300 dilution in 3% BSA/PBS).

**Embryoid body formation**

Pluripotent colonies were dissociated into clumps using EDTA and cultured in DMEM+10%FBS in non-adherent plates for 10 to 14days to form embryoid bodies (EBs). EBs were subsequently plated onto FBS-coated 24 well plates and cultured in DMEM+10%FBS for two weeks. After culture, cells were fixed in 4% paraformaldehyde, and immunofluorescence detection of common differentiation markers of the three germ layers- Ectoderm marker: neurofilament (R&D Systems); Endoderm marker: GATA6 (Cell Signalling Technologies) and Mesoderm marker: Alpha-smooth muscle actin, (R&D Systems) were performed to indicate ability to differentiate into all three germ layers.

**Flow cytometry**

Cells were dissociated using TryPLE™ at 37°C for 5 minutes and counted, then transferred into 1.5 ml Eppendorf tubes. For staining, cells were incubated on ice for 30 minutes in a 5% fetal bovine serum-PBS (FBS-PBS) staining solution containing antibodies to CD90, CD44 CD73 or CD105 (BD Stemflow™ Human MSC Analysis Kit). After incubation, cells were resuspended in 2% FBS-PBS and analysed by flow cytometry using an LSRFortessa™ (BD) and BD FACSDiva™ Software for Windows (Version 8.0). Cells of interest were identified based on their size and granularity through Forward vs Side Scatter (FSC vs SSC) gating. FSC-Height (FSC-H) vs FSC-Area (FSC-A) plots were used to exclude doublets. Unstained cells and single-stained compensation controls were used to set flow cytometer parameters and gating. At least 10,000 events per condition were collected. Data analysis was performed using FlowJo™ Software for Windows (Version 10.6.8.).

**RNA extraction and gene expression analysis by RT-qPCR**

RNA was extracted using QIAGEN RNeasy mini kit according for the manufacturers instructions. For pluripotent and iMSC stages QIAGEN RLT buffer was added directly to cells following PBS wash. For cartilage pellet stages, pellets were washed with PBS and then transferred to 1.5ml tubes containing RLT buffer containing molecular grinding resin. Pellets were then homogenised into the RLT buffer using a plastic pestle, before RNA extraction was followed according to the manufacturer instructions.

RNA was quantified before conversion of 2µg to cDNA with a high capacity cDNA reverse transcription kit (Thermo Fisher Scientific, #4368813. RT-qPCR for gene expression was assessed using gene-specific primers and Power SYBR Green PCR Master Mix (Applied

Biosystems, UK, #4309155) with a Bio-Rad C1000Touch™ Thermal Cycler. Gene expression was normalised to GAPDH, and relative gene expression was calculated using the $2^{-\Delta CT}$ method.

**RNA sequencing and bioinformatic analysis**

Unmapped paired-end sequences from an Illumina HiSeq4000 sequencer were tested by FastQC (http://www.bioinformatics.babraham.ac.uk/projects/fastqc/). Sequence adapters were removed, and reads were quality trimmed using Trimmomatic_0.39 (PMID: 24695404). The reads were mapped against the reference human genome (hg38/GRCh38) and counts per gene were calculated using annotation from GENCODE 39 (http://www.gencodegenes.org/) using STAR_2.7.7a (PMID: 23104886). Normalisation, Principal Components Analysis, and differential expression was calculated with DESeq2_1.36.0 (PMID:25516281). An adjusted P value of <0.1 was used as a threshold for significance. Significant genes with log2FC >0.5 or <-0.5 were used for pathway, Reactome and Gene Ontology enrichment analysis, with the exception of when comparing different stages of the differentiation protocol where changes in gene expression were much greater and log2FC >1 or <-1 were used. Reactome and Gene Ontology enrichment analysis was performed using clusterProfiler in R, terms with an adjusted p-value <0.05 were deemed as significant When affected cartilage pellets expressing high levels of MATN3 were compared to unaffected cartilage pellets also expressing high levels of MATN3, a normalised read count of greater than 1200 for MATN3 was considered as high and a read count less than 1200 was considered as low. This cut off was chosen to ensure that all groups had a minimum of 3 replicates.

**Histological analysis and immunocytochemistry of pellets**

Pellets were washed with PBS and fixed with 4%PFA for 4 hours and then kept for up to 2weeks at 4C in 70% ethanol before processing. Pellets were processed using a tissue processor (Leica, #ASP300. They were then embedded into paraffin and cut into 5µm sections using a microtome (Leica, #RM2255) and mounted on Superfrost Plus Adhesion Microscope Slides

(Epredia™, # 12302108). Sections were initially subjected to consecutive washes with xylene and ethanol for deparaffinization and finished by washes with deionised water. Endogenous peroxidase activity was blocked in 1% Hydrogen Peroxide ($H_2O_2$) for 30 minutes. Antigen retrieval was conducted by incubating in Pepsin Reagent solution (Sigma-Aldrich #R2283) at room temperature for 15 minutes. The samples were then blocked for 1 hour with 10% serum of the secondary antibody species at room temperature. The samples were subsequently incubated with primary antibodies (Supplementary Table), in 10% serum, overnight at 4°C, and then incubated 1 hour with secondary antibodies diluted 1:200 (Supplementary Table). The secondary antibody signal was amplified by incubation with Vectastain Elite ABC Reagent (Vector labs, PK-6100) for 30 minutes. The signal was developed using SIGMAFAST™ 3,3'-Diaminobenzidine (DAB) (Merck, #D4418) for up to 10 minutes at room temperature. Samples were counterstained for 30 seconds in a Mayer's Modified Haematoxylin solution (Abcam, #220365).

**Western blotting**

Samples were washed briefly with 1x PBS, and protein extracted using 1x Cell Lysis buffer with 6μM Phenylmethanesulfonyl Fluoride (CST 9803 and CST 8553) added prior to use. Plates were incubated for 5 minutes on ice, before scraping into eppendorf tubes.  Samples were centrifuged at 14,000xg for 10 minutes at 4°C and supernatant was stored at -20°C until required.

Protein concentration was measured using the Pierce BCA Assay kit (Thermo 23225) and 20μg samples were heated with Pierce lane marker reducing buffer (Thermo 39000) at 95°C for 10 minutes. Samples were run on 10% Bis-Tris gels (Thermo NW00100BOX) with broad range markers (11-245KDa, NEB P7712S). Protein was transferred using the iBlot-2 Gel Transfer device (Thermo IB21001), using iBlot-2 Transfer stacks (nitrocellulose membrane, Thermo IB23001). Membranes were blocked for 1hr at room temperature with 5% BSA in 1x TBS-

0.1%Tween, then probed with the appropriate antibodies and left overnight on a rocker at 4°C. Membranes were washed three times with 1x TBS-0.1%Tween, then secondary antibodies added (1:15,000 IRDye 680RD Donkey [illegible]-Rabbit; LI-COR 925-68071) in 5% BSA in 1x TBS-0.1%Tween. After three washes in 1x TBS-0.1%Tween, staining was imaged and analysed using the Odyssey CLx imaging system (LI-COR) and software.

**Electron Microscopy**

Specimens were fixed with 4% formaldehyde and 2.5% glutaraldehyde in 0.1M Hepes buffer (pH 7.2) overnight. Next day they were poststained with reduced osmium (1% $OsO_4$ and 1.5% $K_4Fe(CN)_6$ in 0.1M Cacodylate buffer, pH 7.2) for 1 hour and after washing with cacodylate buffer they were treated with 1% tannic acid in 0.1M cacodylate buffer for 1 hour. After that they were stained with 1% uranyl acetate in water overnight. Samples were dehydrated with ascending series of ethanol and embedded with TAAB LV epoxy resins. Resulted blocks were cut for 70nm sections with Reichert Ultracut ultramicrotome. Sections were examined with FEI Tecnai 12 Biotwin transmission electron microscope at 100kV and images were taken with Gatan Orius SC1000A bottom mounted CCD camera.

**Cholesterol analysis**

Cholesterol was analysed by gas chromatography (GC) following a modification of the method reported by Barrans et al (17). In brief, cell pellets were homogenized in methanol (1 mL per sample; HPLC grade) in the presence of stigmasterol (internal standard; 20.6 µg per sample); the homogenate was centrifuged (3000 rpm, 4 °C, 5 min) to remove any denatured proteins. The supernatant containing the lipid extract was collected in a clean glass tube and dried under nitrogen. The protein pellet was also collected and stored at -20 °C. The dried lipid extract was reconstituted in ethyl acetate (100 µL; HPLC grade) and analysed on a GC system (6850 Network, Agilent) with flame ionization detector, using a capillary column (Zebron ZB-1, 15 m x 0.32 mm x 0.5 µm; Phenomenex). A calibration line was used to quantify the amount of cholesterol per sample; data were normalized to the amount of sample protein. The protein pellets collected after the extraction of lipids, were used to estimate sample protein as previously reported (18). In brief, protein pellets were solubilized using NaOH (1M) and analysed using a Protein Assay Kit (Bio-Rad, Hercules, CA, USA) as per the manufacturer's instructions.
