## Supplementary figures S1 to S14 and legends for "Human pluripotent stem cell model of multiple epiphyseal dysplasia with *MATN3* mutation identifies altered Matrix organisation and upregulation of the cholesterol biosynthesis pathway"

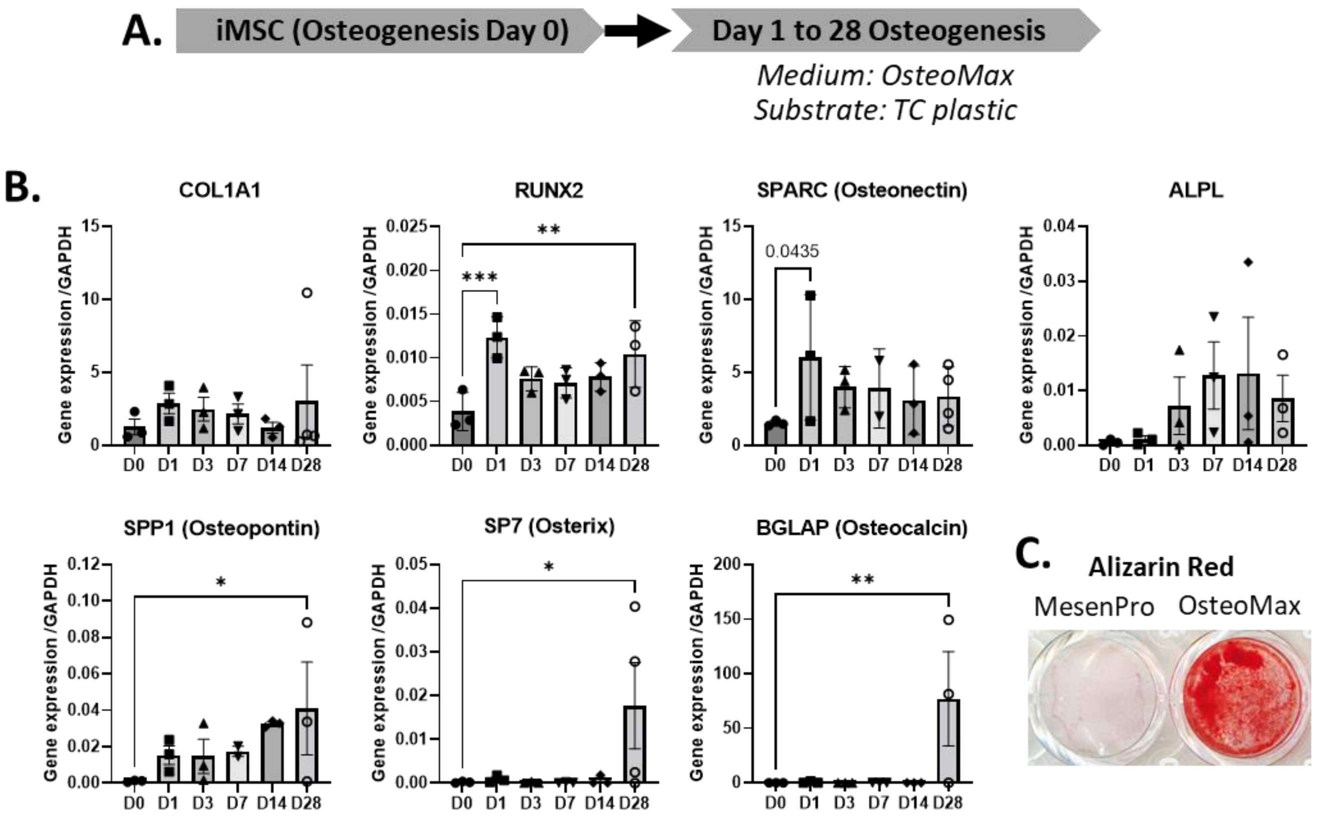


Figure S1. Osteogenic differentiation of pluripotent stem cell derived iMSCs. A, schematic showing osteogenic differentiation protocol for pluripotent stem cell derived iMSCs. B, RT-PCR for osteogenic differentiation markers. C, Alizarin Red staining for mineralisation of D28 osteogenic differentiation (OsteoMax) or control iMSCs grown in mesenPro expansion medium for 28days. Data combined from 3 independent differentiation runs. Statistical differences were calculated using ordinary one-way ANOVA.

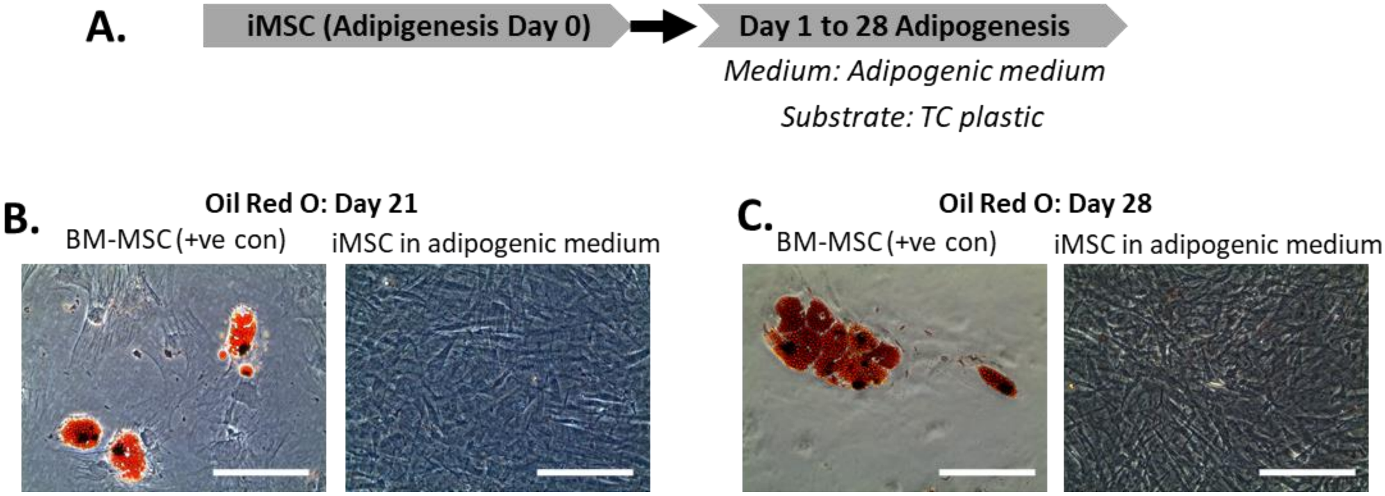


Figure S2. Adipogenic differentiation of pluripotent stem cell derived iMSCs. A, schematic showing adipogenic differentiation protocol for pluripotent stem cell derived iMSCs. B, Oil Red O staining of BM-MSCs (positive control) and iMSCs following 21 days of adipogenic differentiation. C, Oil Red O staining of BM-MSC (positive control) and iMSCs following 28 days of adipogenic differentiation. Scale bars represent 200μm.

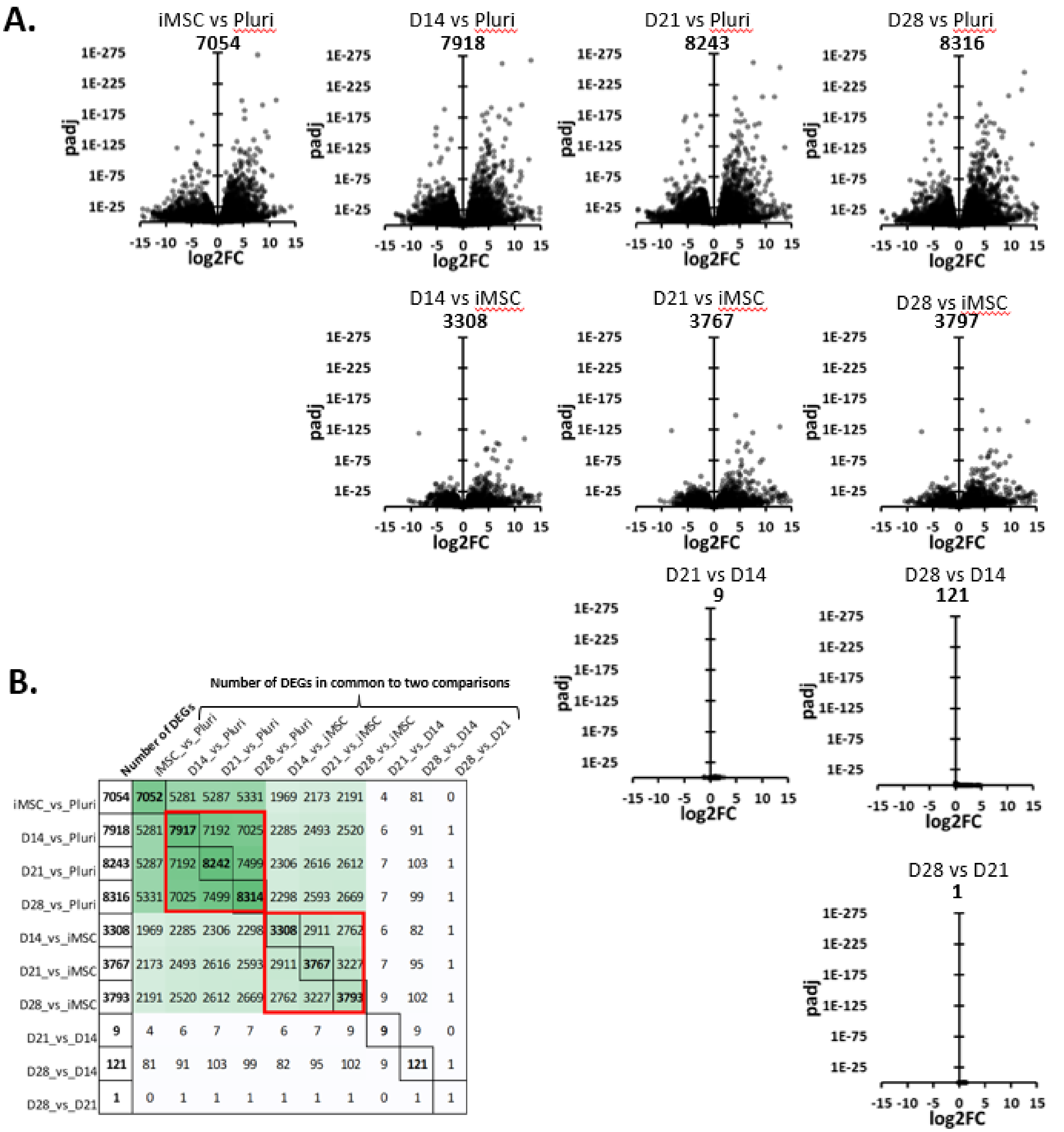


| | Number of DEGs | iMSC_vs_Pluri | D14_vs_Pluri | D21_vs_Pluri | D28_vs_Pluri | D14_vs_iMSC | D21_vs_iMSC | D28_vs_iMSC | D21_vs_D14 | D28_vs_D14 | D28_vs_D21 |
|---|---|---|---|---|---|---|---|---|---|---|---|
| iMSC_vs_Pluri | 7054 | 7052 | 5281 | 5287 | 5331 | 1969 | 2173 | 2191 | 4 | 81 | 0 |
| D14_vs_Pluri | 7918 | 5281 | 7917 | 7192 | 7025 | 2285 | 2493 | 2520 | 6 | 91 | 1 |
| D21_vs_Pluri | 8243 | 5287 | 7192 | 8242 | 7499 | 2306 | 2616 | 2612 | 7 | 103 | 1 |
| D28_vs_Pluri | 8316 | 5331 | 7025 | 7499 | 8314 | 2298 | 2593 | 2669 | 7 | 99 | 1 |
| D14_vs_iMSC | 3308 | 1969 | 2285 | 2306 | 2298 | 3308 | 2911 | 2762 | 6 | 82 | 1 |
| D21_vs_iMSC | 3767 | 2173 | 2493 | 2616 | 2593 | 2911 | 3767 | 3227 | 7 | 95 | 1 |
| D28_vs_iMSC | 3793 | 2191 | 2520 | 2612 | 2669 | 2762 | 3227 | 3793 | 9 | 102 | 1 |
| D21_vs_D14 | 9 | 4 | 6 | 7 | 7 | 6 | 7 | 9 | 9 | 9 | 0 |
| D28_vs_D14 | 121 | 81 | 91 | 103 | 99 | 82 | 95 | 102 | 9 | 121 | 1 |
| D28_vs_D21 | 1 | 0 | 1 | 1 | 1 | 1 | 1 | 1 | 0 | 1 | 1 |

Figure S3. Differential gene expression analysis of unaffected hESCs chondrogenic differentiation. A, Volcano plots comparing each stage of differentiation to another. Numbers above plots indicate the number of DEGs for that comparison. Note the relatively small number of DEGs between D14, D21 and D28 of the cartilage pellet stage. B, Pair-wise analysis of the comparisons in A to find the number of common DEGs between each comparison. Note the high number of common DEGs for each of the D14, D21 and D28 cartilage pellet stages when compared to the earlier pluripotent and iMSC stages.

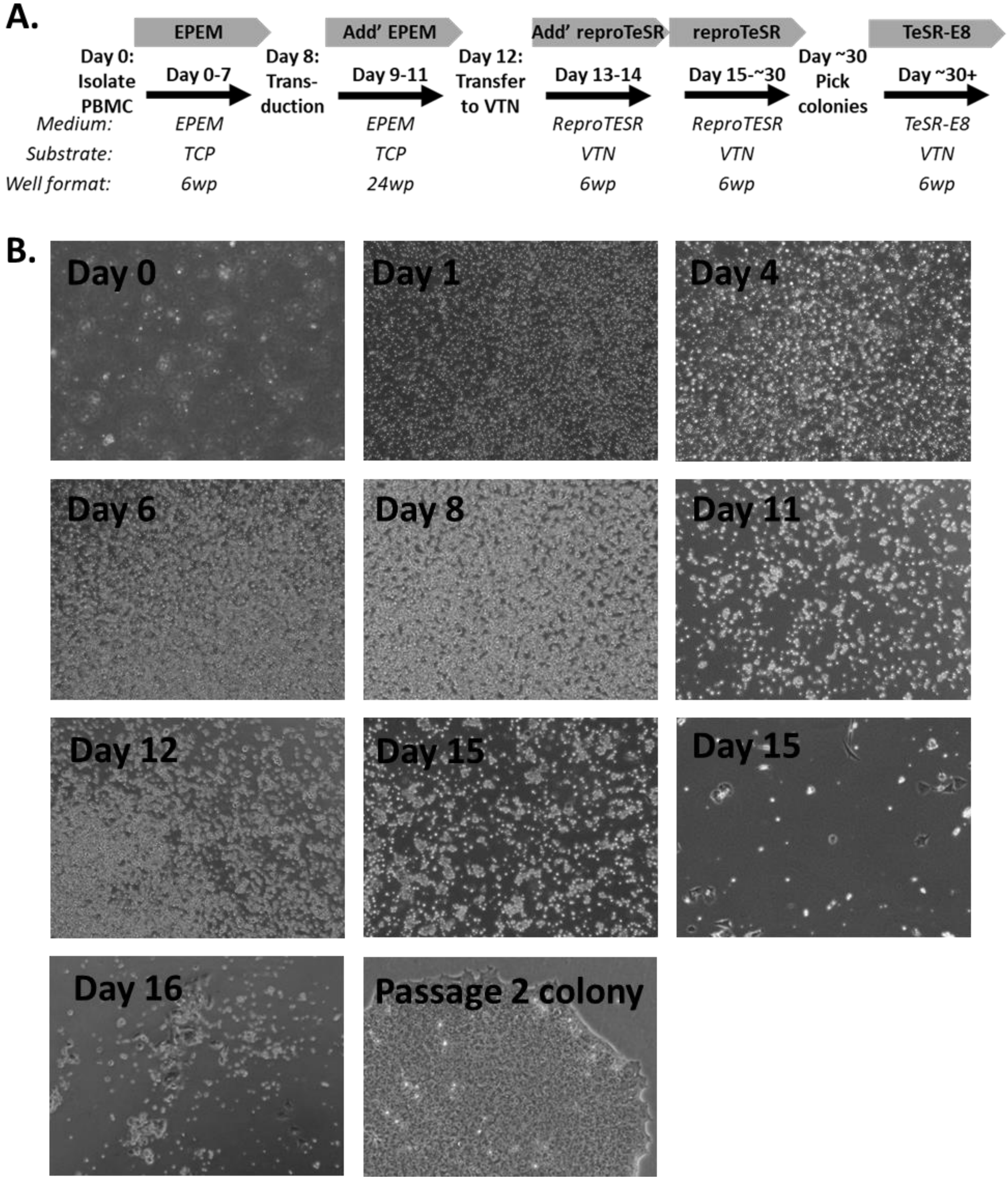


Figure S4. Protocol for reprogramming PBMCs into iPSCs. A, schematic showing reprogramming process, following isolation PBMCs are expanded using erythroid progenitor expansion medium. Reprogramming was performed using CytoTune™-iPS 2.0 Sendai and reproTeSR medium before cells were transferred to TeSR-E8 for expansion. B, Example phase contrast images showing changes in cell morphology during the reprogramming protocol.

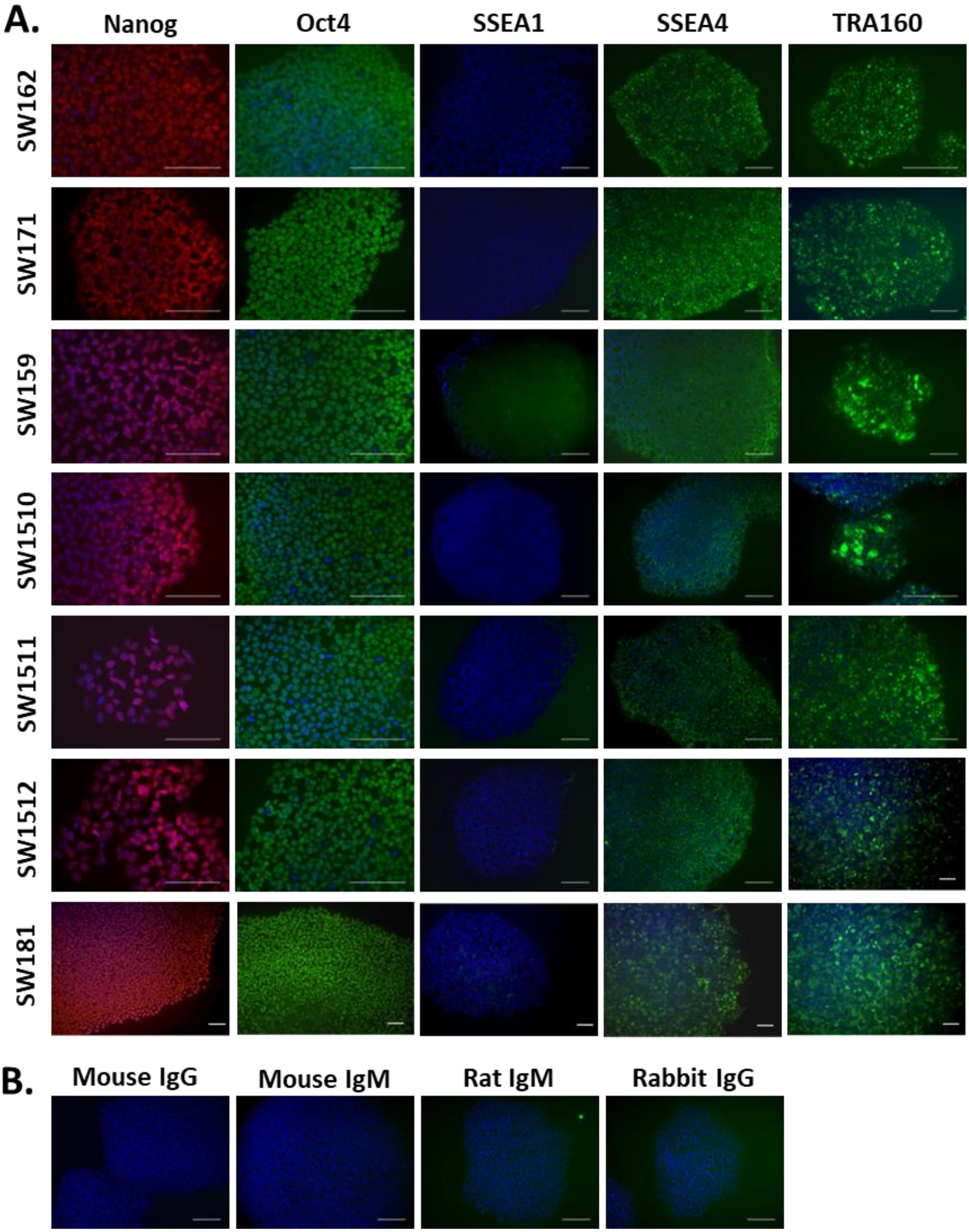


Figure S5. Expression of pluripotency markers in patient derived iPSCs. A, Immunocytochemistry of the pluripotency markers Nanog, Oct4, SSEA1, SSEA4 and TRA160 for the generated unaffected SW162, SW171, SW159 and affected SW1510, SW1511, SW1512, SW181 iPSC lines. B, Isotype controls for antibodies against pluripotency markers.

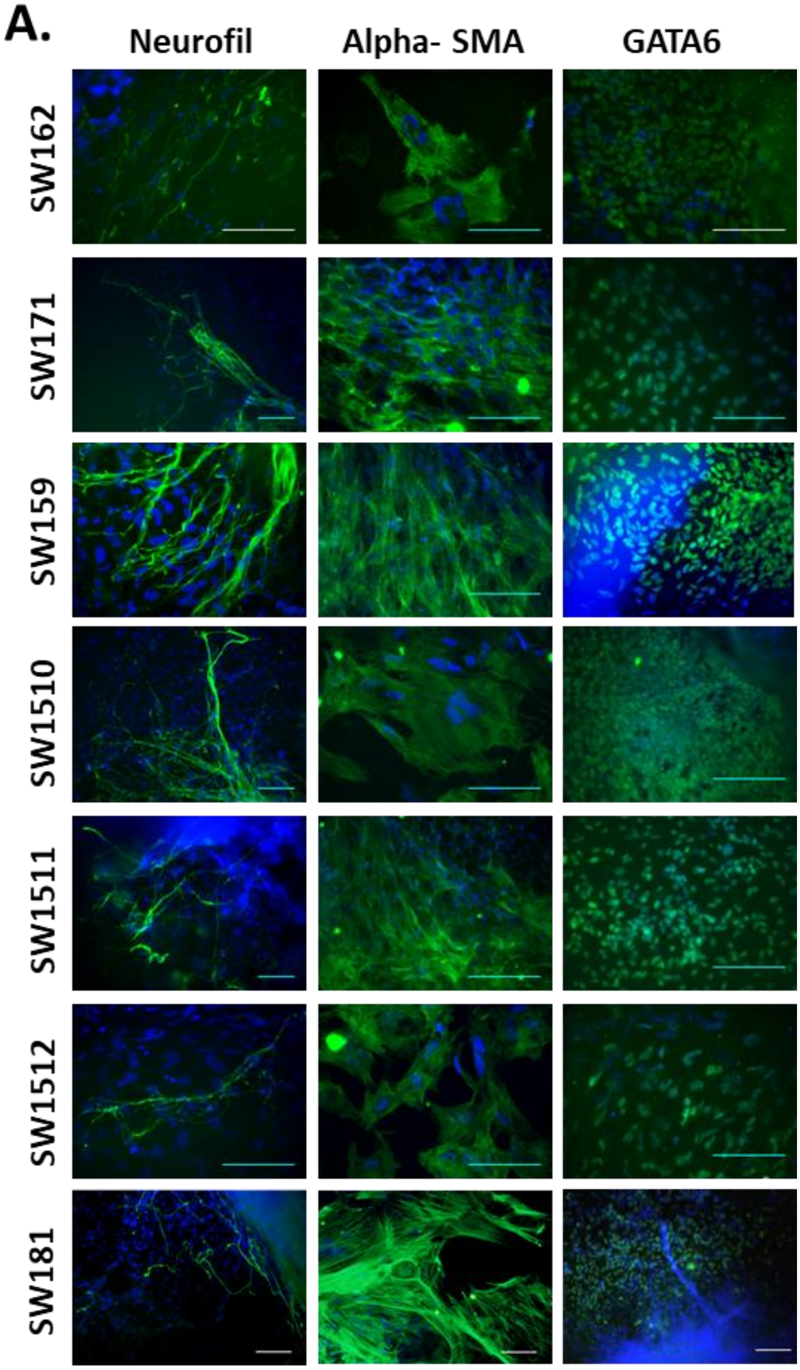


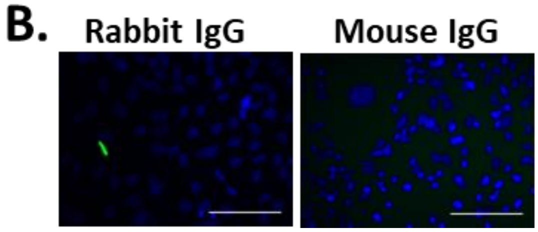


Figure S6. Differentiation of patient derived iPSCs to the three germ layers in EBs. A, Immunocytochemistry for Neurofilament (Neurofil- ectoderm marker), Alpha smooth muscle actin (Alpha-SMA -mesoderm marker) and GATA binding protein 6 (GATA6- endoderm marker) following iPSC differentiation to embryoid bodies (EBs) for unaffected SW162, SW171, SW159 and affected SW1510 (V194D), SW1511 (V194D), SW1512 (V194D), SW181 (T195K) iPSC lines. B, Isotype controls for antibodies against markers of the three germ layers.

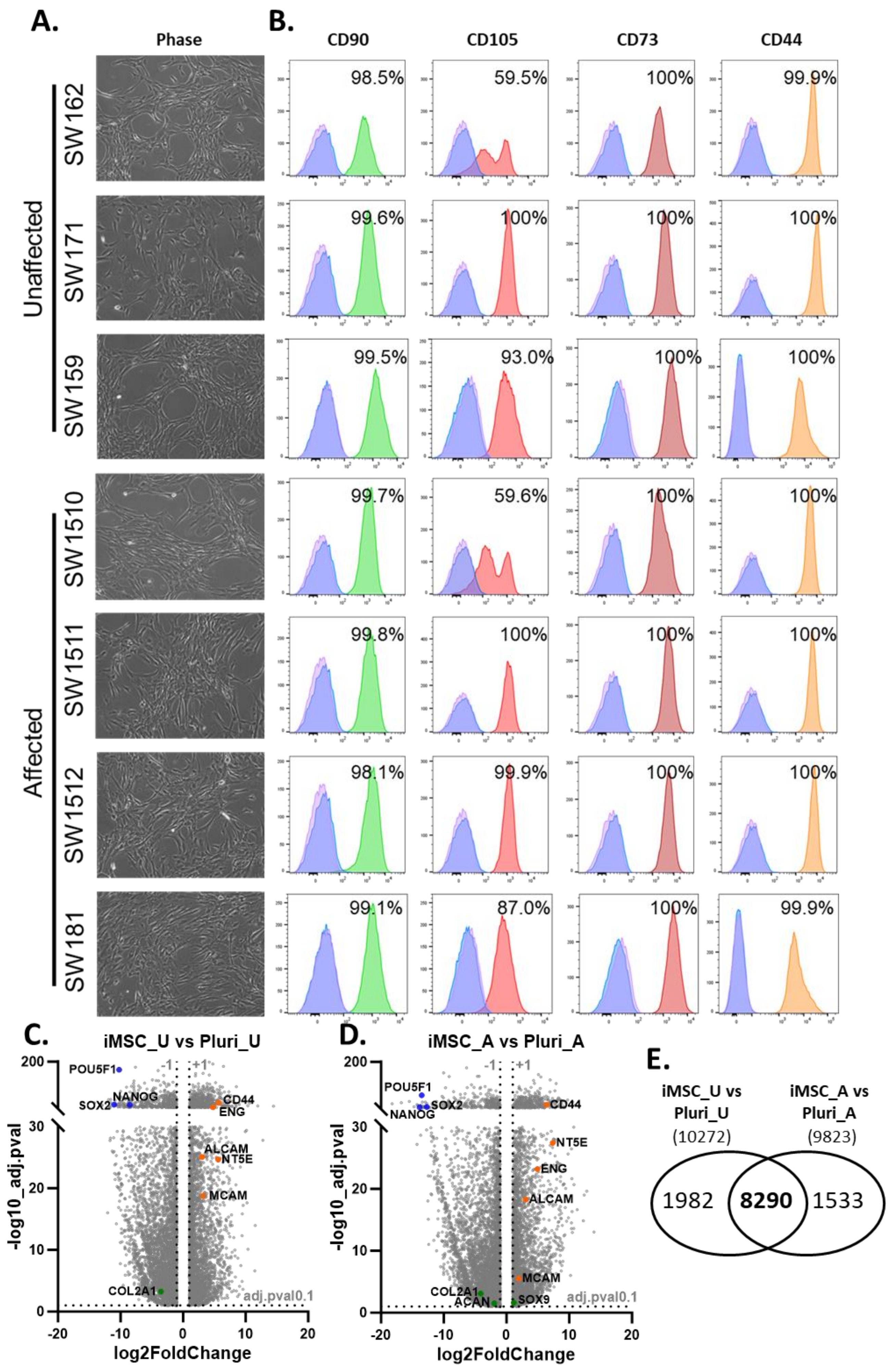
A.
Phase
Unaffected
SW162
SW171
SW159
Affected
SW1510
SW1511
SW1512
SW181
B.
CD90
CD105
CD73
CD44
98.5%
59.5%
100%
99.9%
99.6%
100%
100%
100%
99.5%
93.0%
100%
100%
99.7%
59.6%
100%
100%
99.8%
100%
100%
100%
98.1%
99.9%
100%
100%
99.1%
87.0%
100%
99.9%
C.
iMSC_U vs Pluri_U
POU5F1
NANOG
SOX2
CD44
ENG
ALCAM
NT5E
MCAM
COL2A1
adj.pval0.1
-log10_adj.pval
log2FoldChange
D.
iMSC_A vs Pluri_A
POU5F1
SOX2
NANOG
CD44
NT5E
ENG
ALCAM
MCAM
COL2A1
ACAN
SOX9
adj.pval0.1
-log10_adj.pval
log2FoldChange
E.
iMSC_U vs Pluri_U
(10272)
iMSC_A vs Pluri_A
(9823)
1982
8290
1533

Figure S7. Affected and Unaffected iPSCs can differentiate to iMSCs. A, Phase contrast images of iMSCs generated from unaffected SW162, SW171, SW159 and affected SW1510 (V194D), SW1511 (V194D), SW1512 (V194D), SW181 (T195K) iPSC lines. B, Flow cytometry analysis for CD44, CD73, CD90 and CD105 surface marker expression of iMSCs generated from unaffected and affected iPSC lines. Pink represents non-stained (no antibody), blue represents isotype control antibody and green/red/dark red/yellow represents antibody to marker indicated above. Percentage of positive cells was determined by setting the gating threshold at the 99th percentile of the isotype control fluorescence intensity. C, RNAseq volcano plots showing comparison of iMSC stage with pluripotent stage for unaffected iPSC differentiation. D, RNAseq volcano plots showing comparison of iMSC stage with pluripotent stage for affected iPSC differentiation. Example DEGs associated with; pluripotency are highlighted in blue, mesenchymal cells are highlighted in orange and chondrogenesis are highlighted in green. E, Venn diagrams illustrating the large number of common DEGs between affected and unaffected iPSC differentiation to iMSCs.

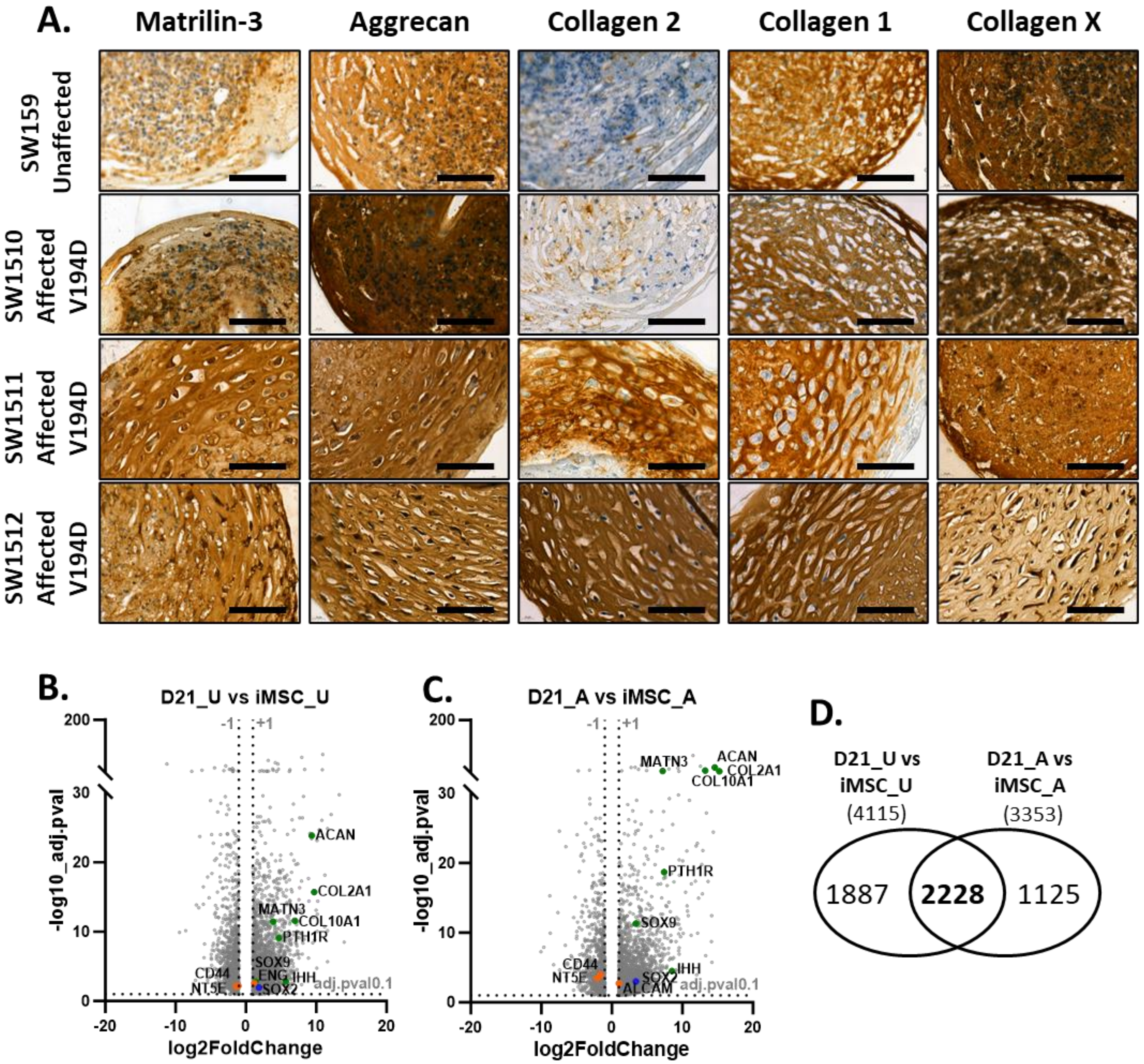


Figure S8. Affected and Unaffected iMSCs can differentiate to chondrocytes. A, Histological analysis of unaffected (SW159) and affected (SW1510, SW1511, SW1512) chondrogenesis at day 21 of pellet culture. Scale bars = 100µm. B, RNAseq volcano plots showing comparison of D21 cartilage pellets with iMSC stage for unaffected and affected iPSC differentiation. Example DEGs associated with; pluripotency are highlighted in blue, mesenchymal cells are highlighted in orange and chondrogenesis are highlighted in green. D, Venn diagrams illustrating the large number of common DEGs between affected and unaffected differentiation of iMSCs to cartilage pellets.

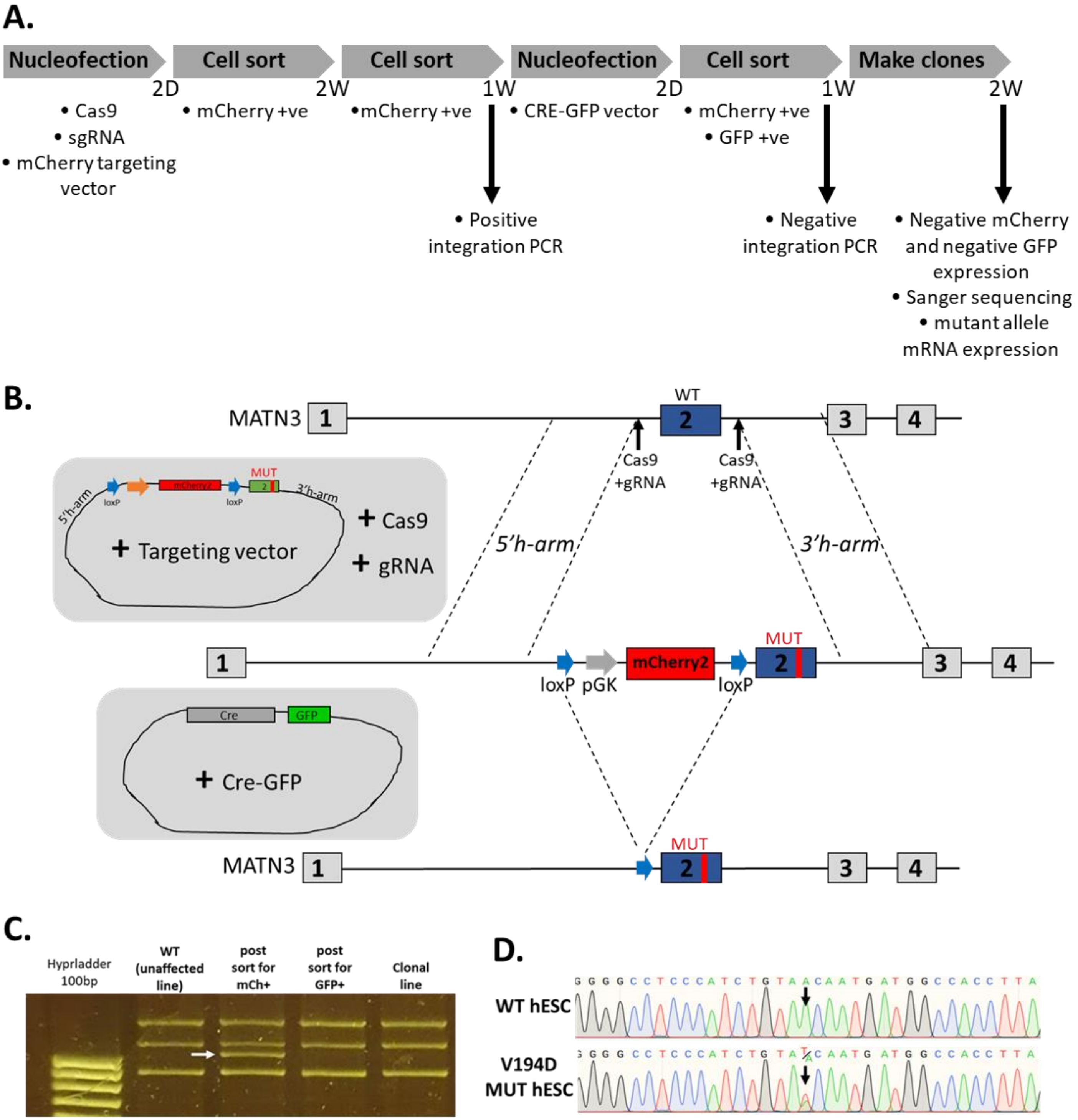
A.
Nucleofection
Cell sort
Cell sort
Nucleofection
Cell sort
Make clones
2D
2W
1W
2D
1W
2W
• Cas9
• sgRNA
• mCherry targeting vector
• mCherry +ve
•mCherry +ve
• CRE-GFP vector
• mCherry +ve
• GFP +ve
• Positive integration PCR
• Negative integration PCR
• Negative mCherry and negative GFP expression
• Sanger sequencing
• mutant allele mRNA expression
B.
MATN3
WT
Cas9 +gRNA
Cas9 +gRNA
5'h-arm
3'h-arm
MUT
loxP
loxP
+ Targeting vector
+ Cas9
+ gRNA
mCherry2
loxP
pGK
MUT
Cre
GFP
+ Cre-GFP
MATN3
MUT
C.
Hyprladder 100bp
WT (unaffected line)
post sort for mCh+
post sort for GFP+
Clonal line
D.
WT hESC
V194D MUT hESC

Figure S9. Generation of heterozygous MATN3 V194D mutation in hESC using CRISPR-Cas9. A, schematic illustrating the timeline of the method used to create clonal hESC lines with heterozygous MATN3 V194D mutation. Following nucleofection with Cas9, sgRNA and mCherry targeting vector. mCherry positive cells were first sorted for those that have taken up the vector and then again after two weeks for those that have stable integration of mCherry. Cells were then nucleofected with CRE-recombinase GFP to excise the mCherry selection. They were sorted for mCherry positive and GFP positive cells two days post nucleofection. This third sort and further enrichment for mCherry is possible as it takes longer than two days post CRE-recombinase for mCherry expression to be lost. After additional culture cells became negative for both mCherry and GFP. Following PCR validation clonal lines were selected. B, schematic illustrating the strategy to create heterozygous MATN3 V194D mutation in hESC using CRISPR-Cas9. Cas9:sgRNA targets within the introns surrounding exon 2. The mCherry targeting vector with homology arms against the surrounding sequences then directs homology directed repair (HDR) to replace WT exon 2 with a V194D mutant exon 2 along with mCherry2 driven by pGK surrounded by loxP site within upstream intron. Cre-recombinase then targets the loxP sites to excise mCherry and pGK, leaving behind V194D mutant exon 2 and a loxP scar in the intron. C, PCR showing successful integration and then successful excision of targeting vector. Note there are some non-specific PCR products. D, Sanger sequencing validation of the presence of mutation in CRISPR edited cells.

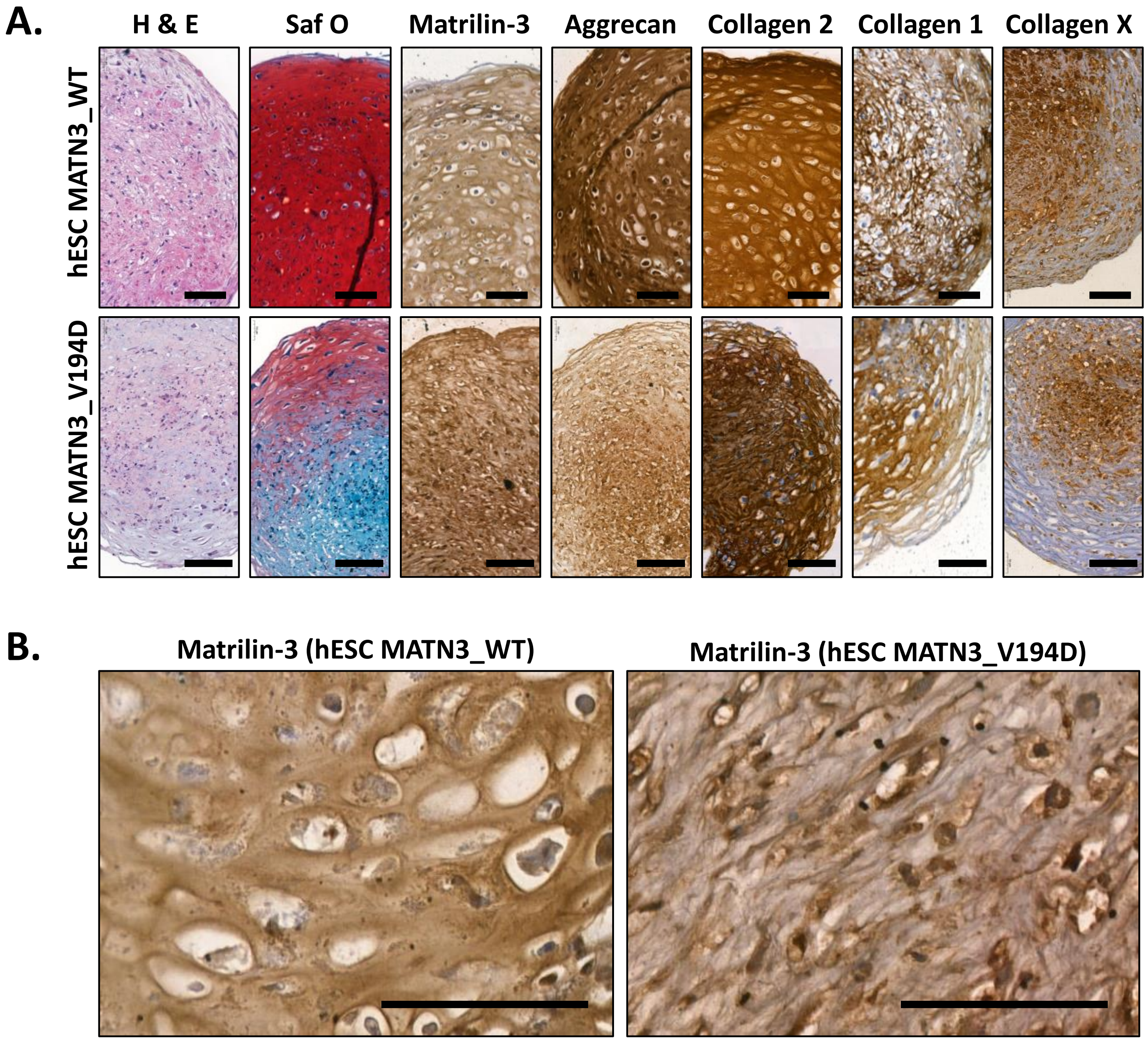


Figure S10. A, Histological analysis of WT and CRISPR-Cas9 V194D MATN3 mutant hESC chondrogenesis. Day 21 cartilage pellets from either WT or CRISPR-Cas9 V194D MATN3 mutant hESCs. B, Higher power images of matrilin-3, note the reduced matrix expression in V194D MATN3 mutant. Scale bars = 100µm.

A.

WT

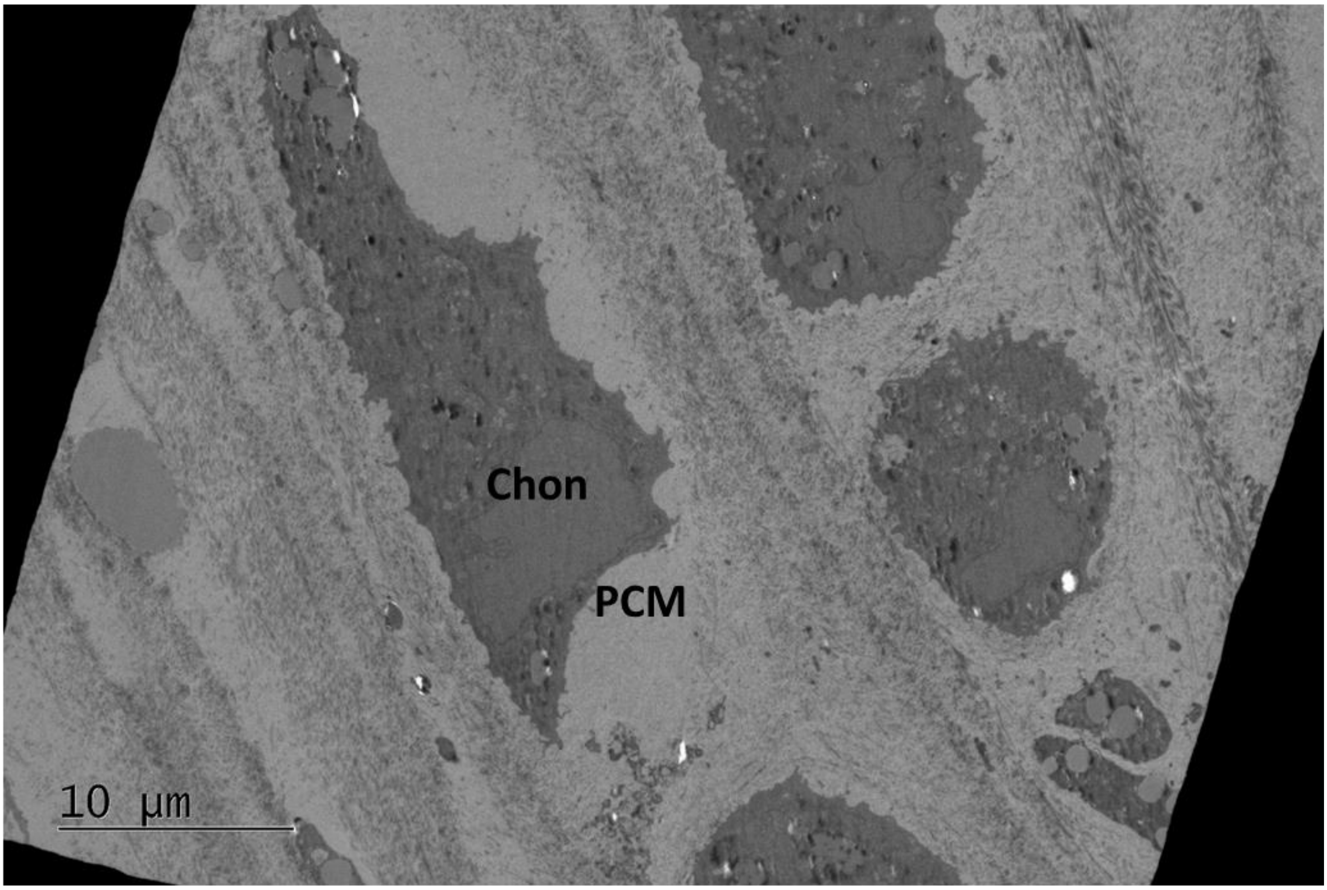


B.

V194D

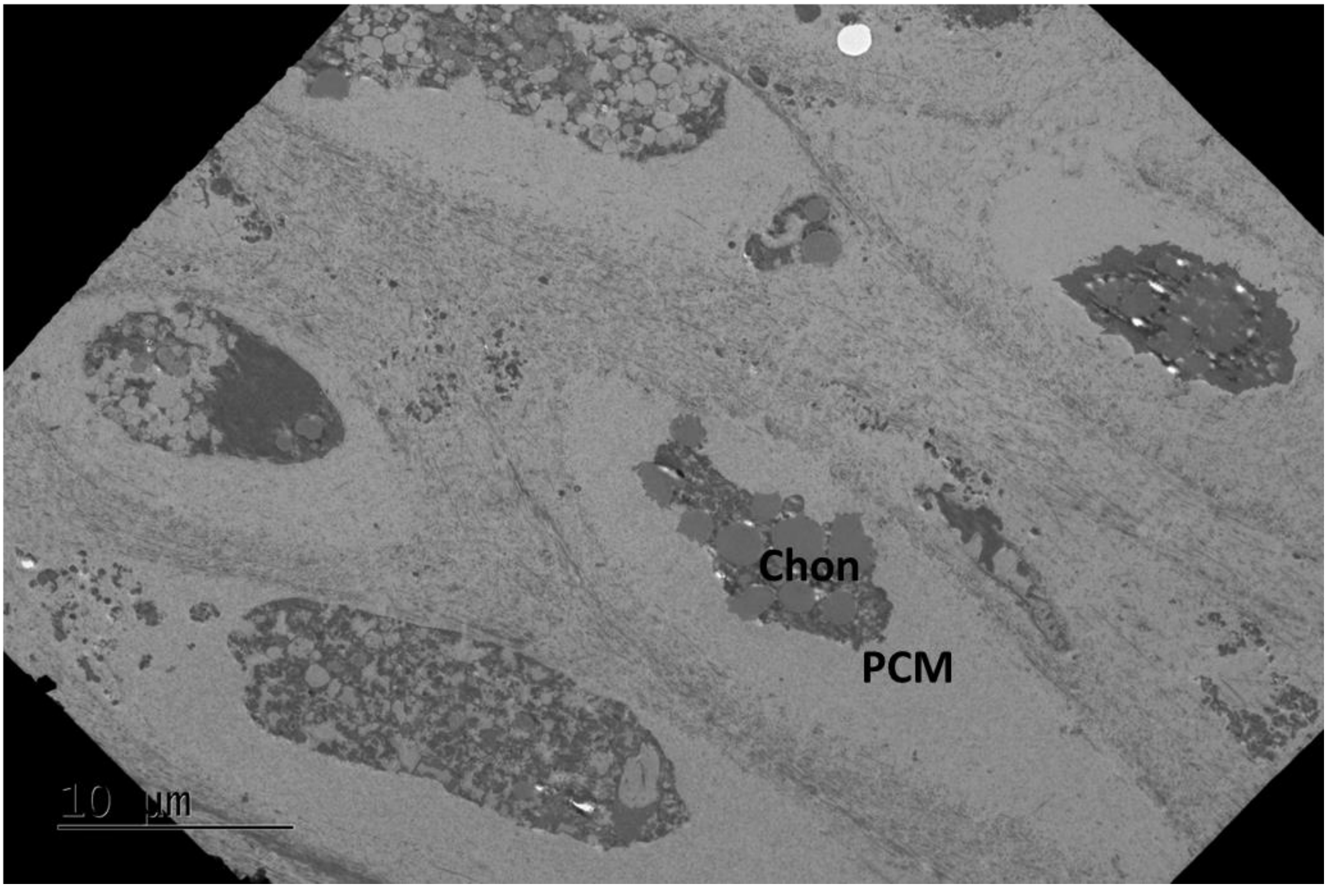

C.

WT

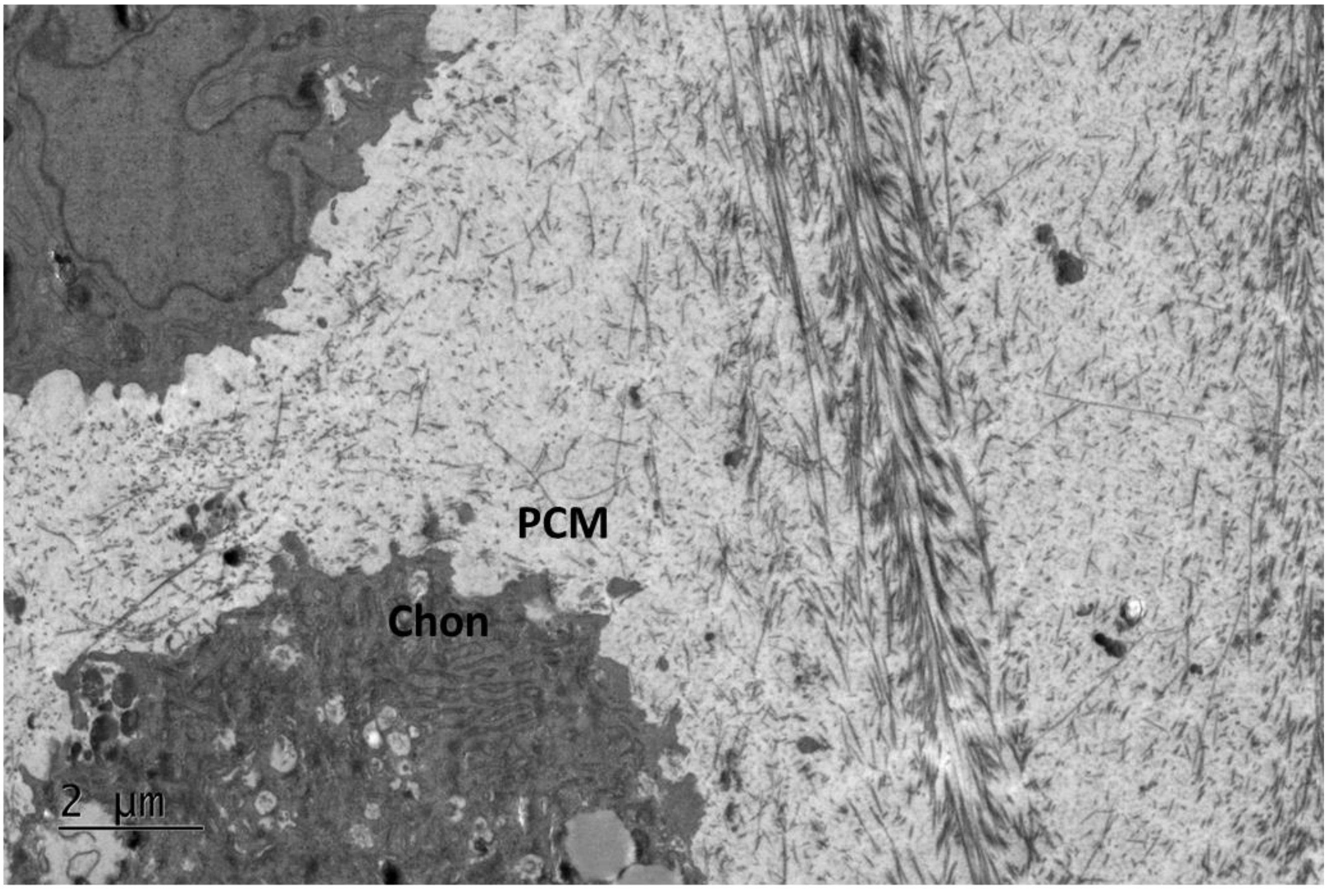


D.

V194D

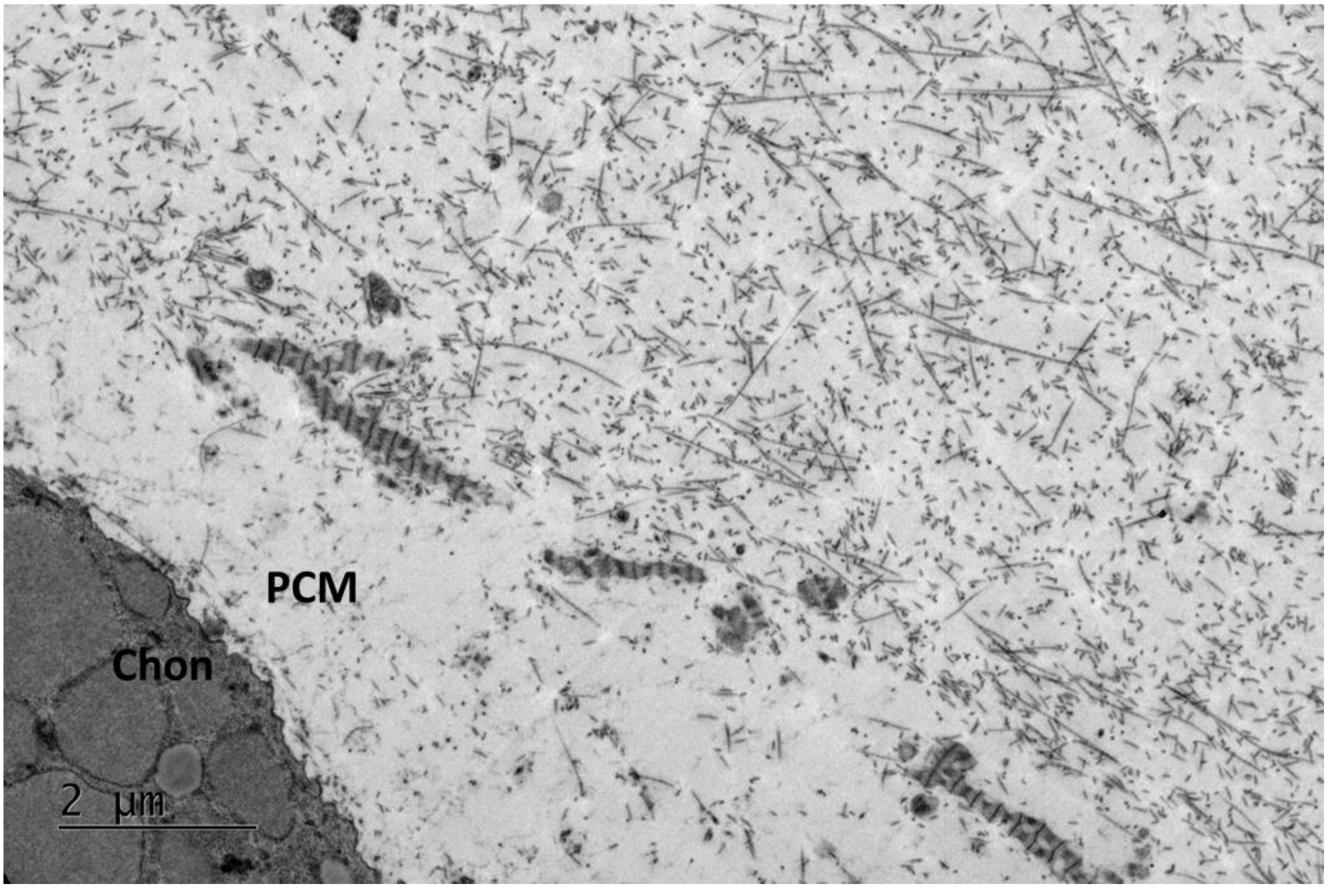

E. WT

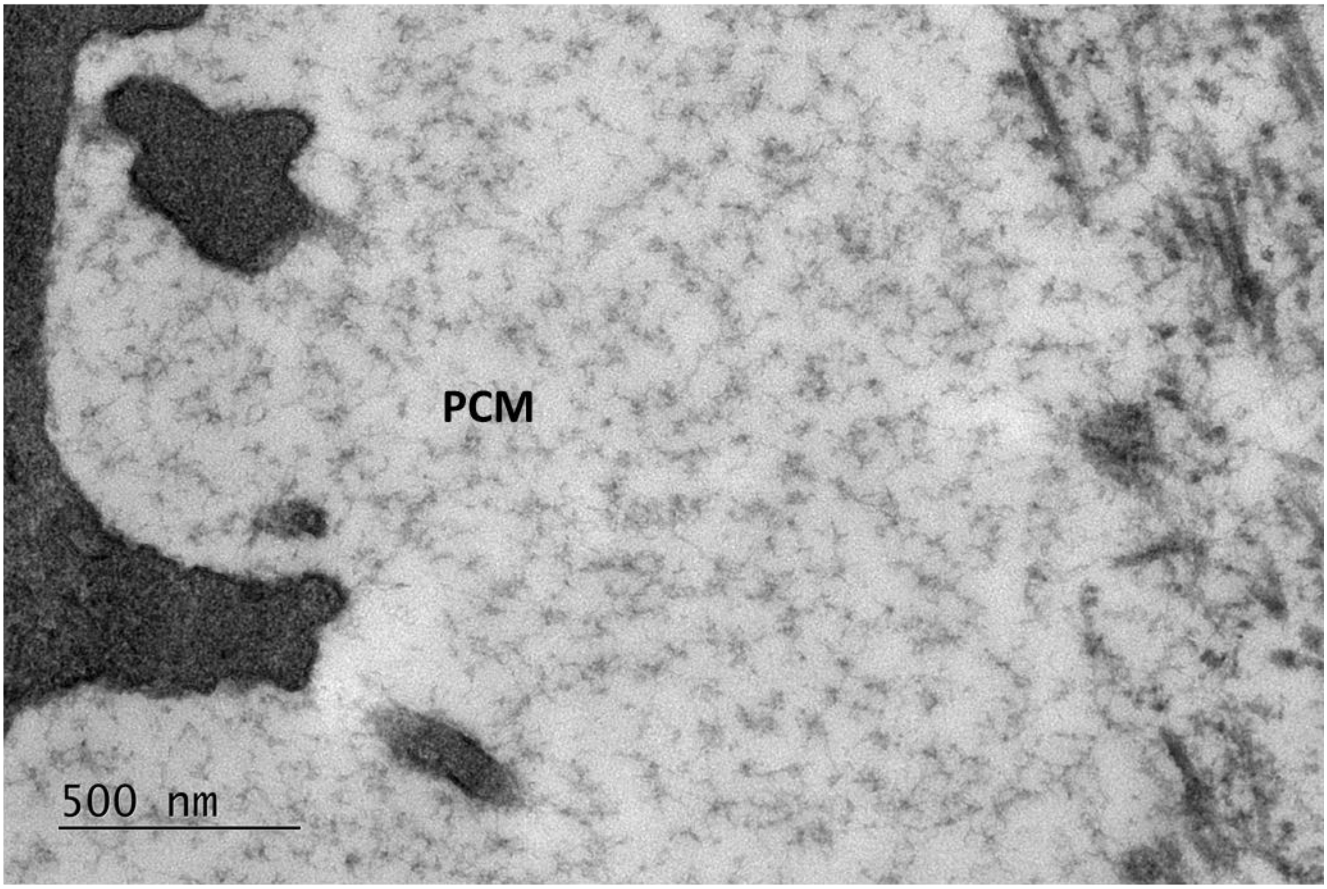
PCM
500 nm

F. V194D

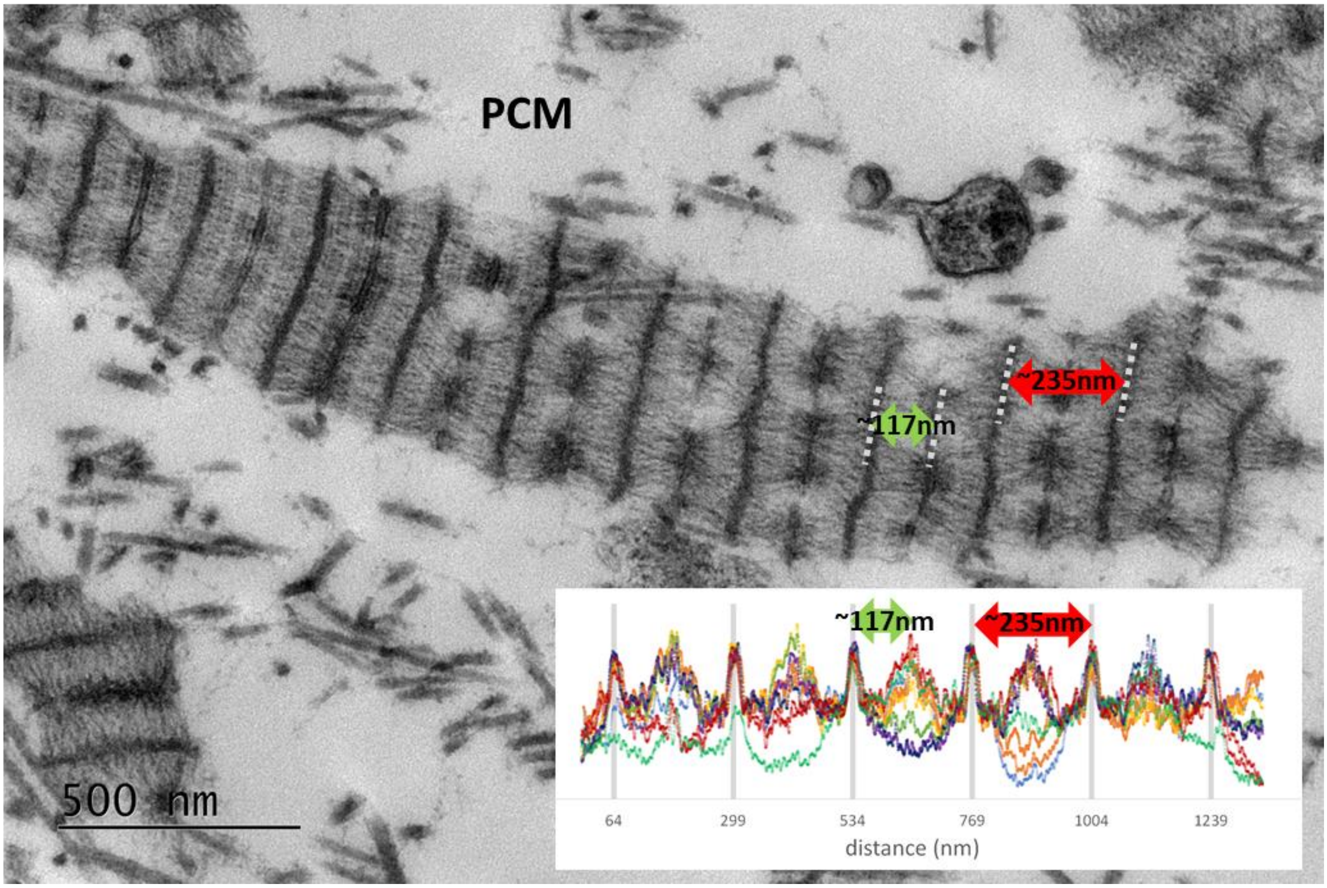
PCM
~117nm
~235nm
~117nm
~235nm
64
299
534
769
1004
1239
distance (nm)
500 nm

**G.** **WT**

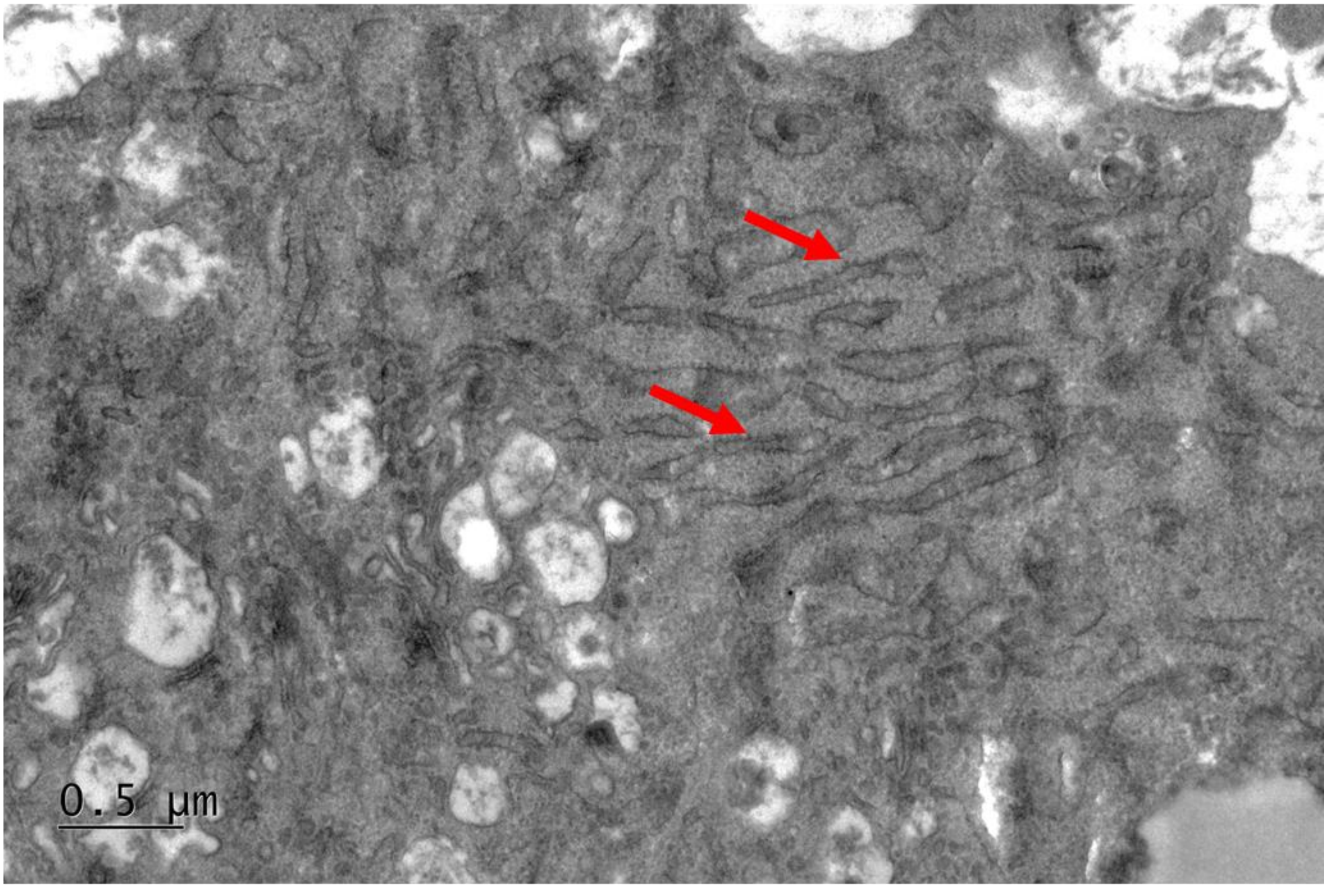


**H.** **V194D**

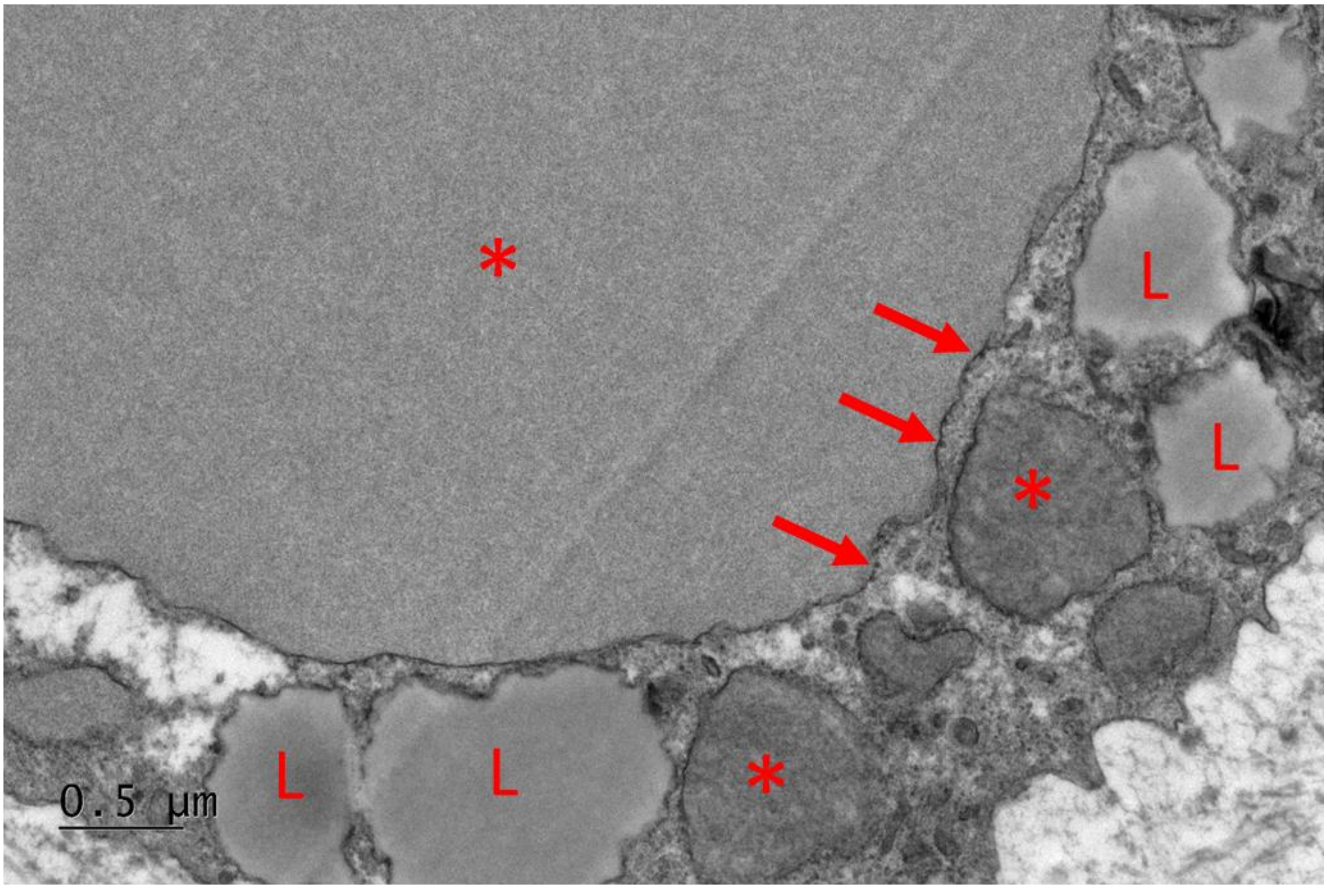

Figure S11. V194D MATN3 mutant cartilage pellet TEM. A and B, zoomed out illustrating the location of chondroncytes (Chon) and their pericellular matrix (PCM) within the WT and V194D mutant cartilage pellets. C and D, same as A and B but at higher magnification. E, pericellular matrix (PCM) region of WT pellets. F, altered matrix structure observed in pericellular matrix (PCM) region of V194D mutant cartilage pellets. Measurement of repeating pattern in matrix structures of V194D mutant cartilage pellets. Distance between darkest bands is ~235nm and distance between darkest band a lighter band is ~117nm. G, Presence of ribosomes on normal endoplasmic reticulum in WT cartilage pellets  H, Presence of ribosomes surrounding distended endoplasmic reticulum in V194D MATN3 mutant cartilage pellets indicated by arrows. ‘*’ indicates presence of distended ER, ‘L’ indicates lipid droplet accumulation.

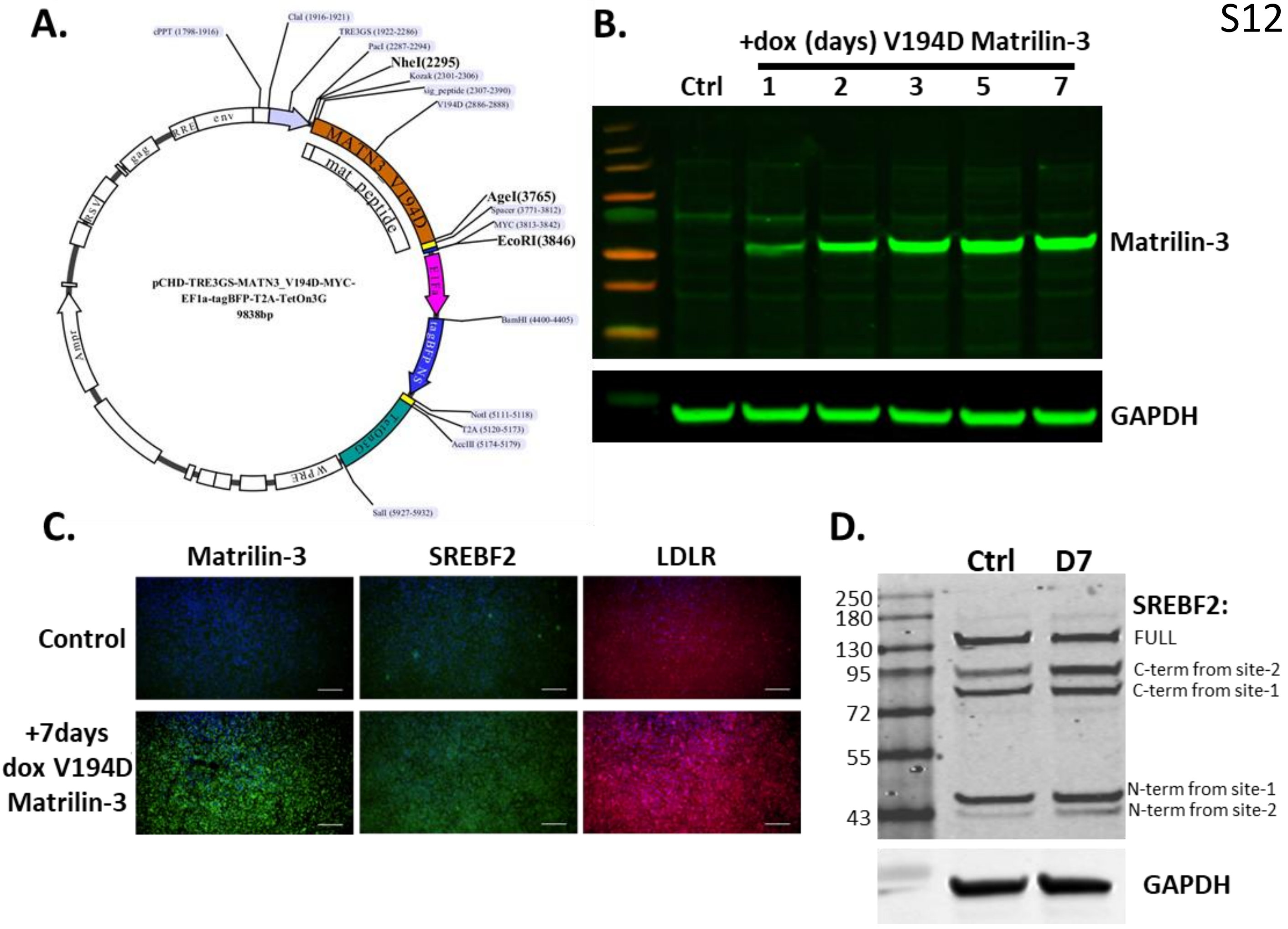


Figure S12. Doxycycline inducible mutant Matrlin-3 in TC28a2 cells causes SREBF2 cleavage and increased expression of LDLR (low-density lipoprotein receptor). A, map of doxycycline inducible V194D mutant matrilin-3 expression vector used to generate TC28a2-dox-MATN3-V194D cell line. B, matrilin-3 Western blot of TC28a2-dox-MATN3-V194D cells following additional of doxycycline for 1 to 7 days. C, immunocytochemistry of matrilin-3, SREBF2 and LDLR in TC28a2-dox-MATN3-V194D cells following addition of doxycycline for 7 days. D, SREBP2 western blot of TC28a2-dox-MATN3-V194D cells following additional of doxycycline for 7 days compared with no doxycycline control.

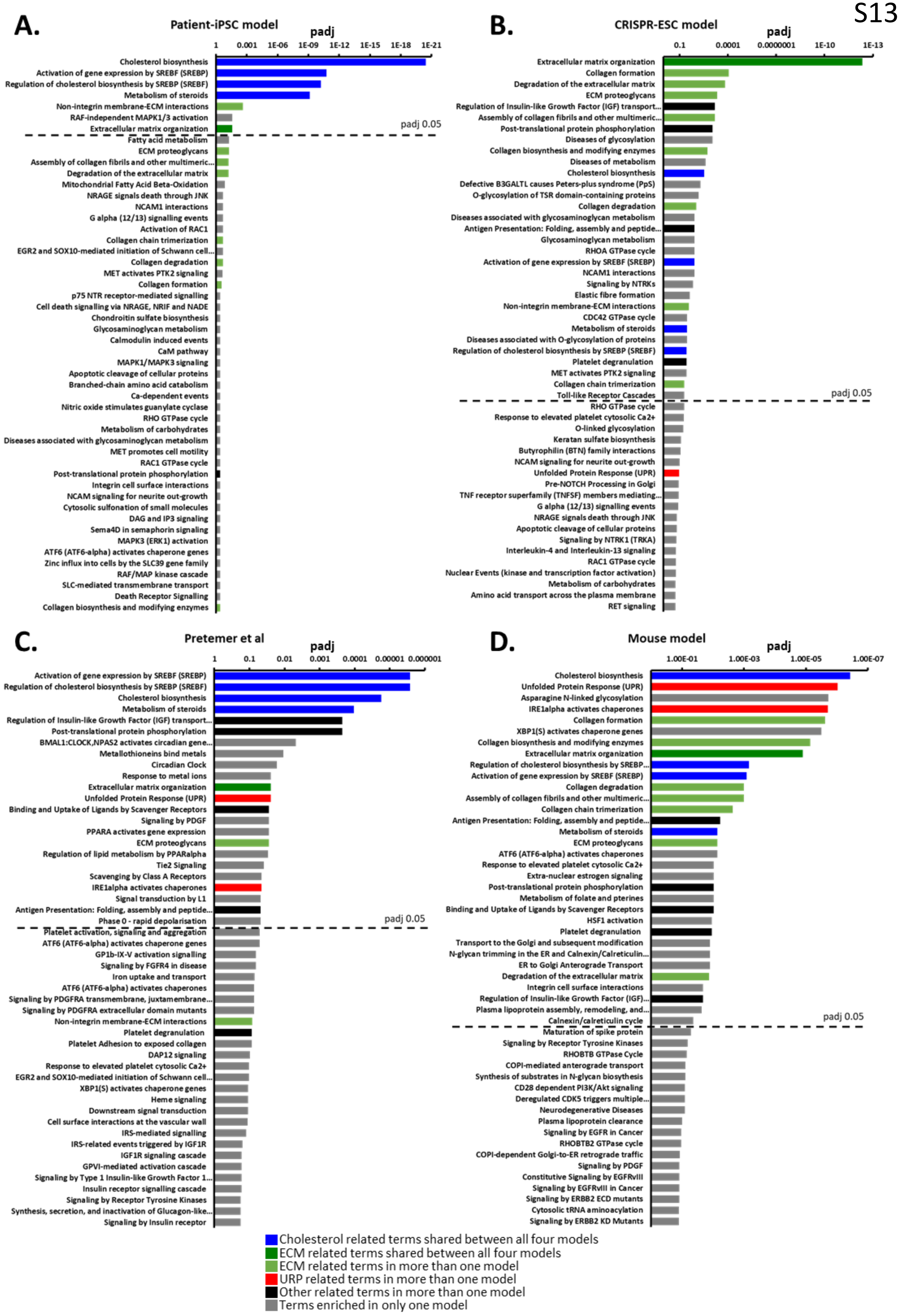

A.
Patient-iPSC model
padj
1
0.001
1E-06
1E-09
1E-12
1E-15
1E-18
1E-21
Cholesterol biosynthesis
Activation of gene expression by SREBF (SREBP)
Regulation of cholesterol biosynthesis by SREBP (SREBF)
Metabolism of steroids
Non-integrin membrane-ECM interactions
RAF-independent MAPK1/3 activation
Extracellular matrix organization
padj 0.05
Fatty acid metabolism
ECM proteoglycans
Assembly of collagen fibrils and other multimeric...
Degradation of the extracellular matrix
Mitochondrial Fatty Acid Beta-Oxidation
NRAGE signals death through JNK
NCAM1 interactions
G alpha (12/13) signalling events
Activation of RAC1
Collagen chain trimerization
EGR2 and SOX10-mediated initiation of Schwann cell...
Collagen degradation
MET activates PTK2 signaling
Collagen formation
p75 NTR receptor-mediated signalling
Cell death signalling via NRAGE, NRIF and NADE
Chondroitin sulfate biosynthesis
Glycosaminoglycan metabolism
Calmodulin induced events
CaM pathway
MAPK1/MAPK3 signaling
Apoptotic cleavage of cellular proteins
Branched-chain amino acid catabolism
Ca-dependent events
Nitric oxide stimulates guanylate cyclase
RHO GTPase cycle
Metabolism of carbohydrates
Diseases associated with glycosaminoglycan metabolism
MET promotes cell motility
RAC1 GTPase cycle
Post-translational protein phosphorylation
Integrin cell surface interactions
NCAM signaling for neurite out-growth
Cytosolic sulfonation of small molecules
DAG and IP3 signaling
Sema4D in semaphorin signaling
MAPK3 (ERK1) activation
ATF6 (ATF6-alpha) activates chaperone genes
Zinc influx into cells by the SLC39 gene family
RAF/MAP kinase cascade
SLC-mediated transmembrane transport
Death Receptor Signalling
Collagen biosynthesis and modifying enzymes
B.
CRISPR-ESC model
padj
0.1
0.0001
0.0000001
1E-10
1E-13
Extracellular matrix organization
Collagen formation
Degradation of the extracellular matrix
ECM proteoglycans
Regulation of Insulin-like Growth Factor (IGF) transport...
Assembly of collagen fibrils and other multimeric...
Post-translational protein phosphorylation
Diseases of glycosylation
Collagen biosynthesis and modifying enzymes
Diseases of metabolism
Cholesterol biosynthesis
Defective B3GALTL causes Peters-plus syndrome (PpS)
O-glycosylation of TSR domain-containing proteins
Collagen degradation
Diseases associated with glycosaminoglycan metabolism
Antigen Presentation: Folding, assembly and peptide...
Glycosaminoglycan metabolism
RHOA GTPase cycle
Activation of gene expression by SREBF (SREBP)
NCAM1 interactions
Signaling by NTRKs
Elastic fibre formation
Non-integrin membrane-ECM interactions
CDC42 GTPase cycle
Metabolism of steroids
Diseases associated with O-glycosylation of proteins
Regulation of cholesterol biosynthesis by SREBP (SREBF)
Platelet degranulation
MET activates PTK2 signaling
Collagen chain trimerization
Toll-like Receptor Cascades
padj 0.05
RHO GTPase cycle
Response to elevated platelet cytosolic Ca2+
O-linked glycosylation
Keratan sulfate biosynthesis
Butyrophilin (BTN) family interactions
NCAM signaling for neurite out-growth
Unfolded Protein Response (UPR)
Pre-NOTCH Processing in Golgi
TNF receptor superfamily (TNFSF) members mediating...
G alpha (12/13) signalling events
NRAGE signals death through JNK
Apoptotic cleavage of cellular proteins
Signaling by NTRK1 (TRKA)
Interleukin-4 and Interleukin-13 signaling
RAC1 GTPase cycle
Nuclear Events (kinase and transcription factor activation)
Metabolism of carbohydrates
Amino acid transport across the plasma membrane
RET signaling
C.
Pretemer et al
padj
1
0.1
0.01
0.001
0.0001
0.00001
0.000001
Activation of gene expression by SREBF (SREBP)
Regulation of cholesterol biosynthesis by SREBP (SREBF)
Cholesterol biosynthesis
Metabolism of steroids
Regulation of Insulin-like Growth Factor (IGF) transport...
Post-translational protein phosphorylation
BMAL1:CLOCK,NPAS2 activates circadian gene...
Metallothioneins bind metals
Circadian Clock
Response to metal ions
Extracellular matrix organization
Unfolded Protein Response (UPR)
Binding and Uptake of Ligands by Scavenger Receptors
Signaling by PDGF
PPARA activates gene expression
ECM proteoglycans
Regulation of lipid metabolism by PPARalpha
Tie2 Signaling
Scavenging by Class A Receptors
IRE1alpha activates chaperones
Signal transduction by L1
Antigen Presentation: Folding, assembly and peptide...
Phase 0 - rapid depolarisation
padj 0.05
Platelet activation, signaling and aggregation
ATF6 (ATF6-alpha) activates chaperone genes
GP1b-IX-V activation signalling
Signaling by FGFR4 in disease
Iron uptake and transport
ATF6 (ATF6-alpha) activates chaperones
Signaling by PDGFRA transmembrane, juxtamembrane...
Signaling by PDGFRA extracellular domain mutants
Non-integrin membrane-ECM interactions
Platelet degranulation
Platelet Adhesion to exposed collagen
DAP12 signaling
Response to elevated platelet cytosolic Ca2+
EGR2 and SOX10-mediated initiation of Schwann cell...
XBP1(S) activates chaperone genes
Heme signaling
Downstream signal transduction
Cell surface interactions at the vascular wall
IRS-mediated signalling
IRS-related events triggered by IGF1R
IGF1R signaling cascade
GPVI-mediated activation cascade
Signaling by Type 1 Insulin-like Growth Factor 1...
Insulin receptor signalling cascade
Signaling by Receptor Tyrosine Kinases
Synthesis, secretion, and inactivation of Glucagon-like...
Signaling by Insulin receptor
D.
Mouse model
padj
1.00E-01
1.00E-03
1.00E-05
1.00E-07
Cholesterol biosynthesis
Unfolded Protein Response (UPR)
Asparagine N-linked glycosylation
IRE1alpha activates chaperones
Collagen formation
XBP1(S) activates chaperone genes
Collagen biosynthesis and modifying enzymes
Extracellular matrix organization
Regulation of cholesterol biosynthesis by SREBP...
Activation of gene expression by SREBF (SREBP)
Collagen degradation
Assembly of collagen fibrils and other multimeric...
Collagen chain trimerization
Antigen Presentation: Folding, assembly and peptide...
Metabolism of steroids
ECM proteoglycans
ATF6 (ATF6-alpha) activates chaperones
Response to elevated platelet cytosolic Ca2+
Extra-nuclear estrogen signaling
Post-translational protein phosphorylation
Metabolism of folate and pterines
Binding and Uptake of Ligands by Scavenger Receptors
HSF1 activation
Platelet degranulation
Transport to the Golgi and subsequent modification
N-glycan trimming in the ER and Calnexin/Calreticulin...
ER to Golgi Anterograde Transport
Degradation of the extracellular matrix
Integrin cell surface interactions
Regulation of Insulin-like Growth Factor (IGF)...
Plasma lipoprotein assembly, remodeling, and...
Calnexin/calreticulin cycle
padj 0.05
Maturation of spike protein
Signaling by Receptor Tyrosine Kinases
RHOBTB GTPase Cycle
COPI-mediated anterograde transport
Synthesis of substrates in N-glycan biosynthesis
CD28 dependent PI3K/Akt signaling
Deregulated CDK5 triggers multiple...
Neurodegenerative Diseases
Plasma lipoprotein clearance
Signaling by EGFR in Cancer
RHOBTB2 GTPase cycle
COPI-dependent Golgi-to-ER retrograde traffic
Signaling by PDGF
Constitutive Signaling by EGFRvIII
Signaling by EGFRvIII in Cancer
Signaling by ERBB2 ECD mutants
Cytosolic tRNA aminoacylation
Signaling by ERBB2 KD Mutants
Cholesterol related terms shared between all four models
ECM related terms shared between all four models
ECM related terms in more than one model
URP related terms in more than one model
Other related terms in more than one model
Terms enriched in only one model

Figure S13. Reactome analysis of patient-iPSC model, CRISPR-hESC model, Pretemer et al model and mouse model. A, 480 DEGs were used an input for patient hiPSC model. B, 1978 DEGs were used as input for CRISPR hESC model. C, 1750 DEGs were used as input for Pretermer et al model . D, 428 DEGs were used as input for mouse model.

**A.** **Patient-iPSC model**

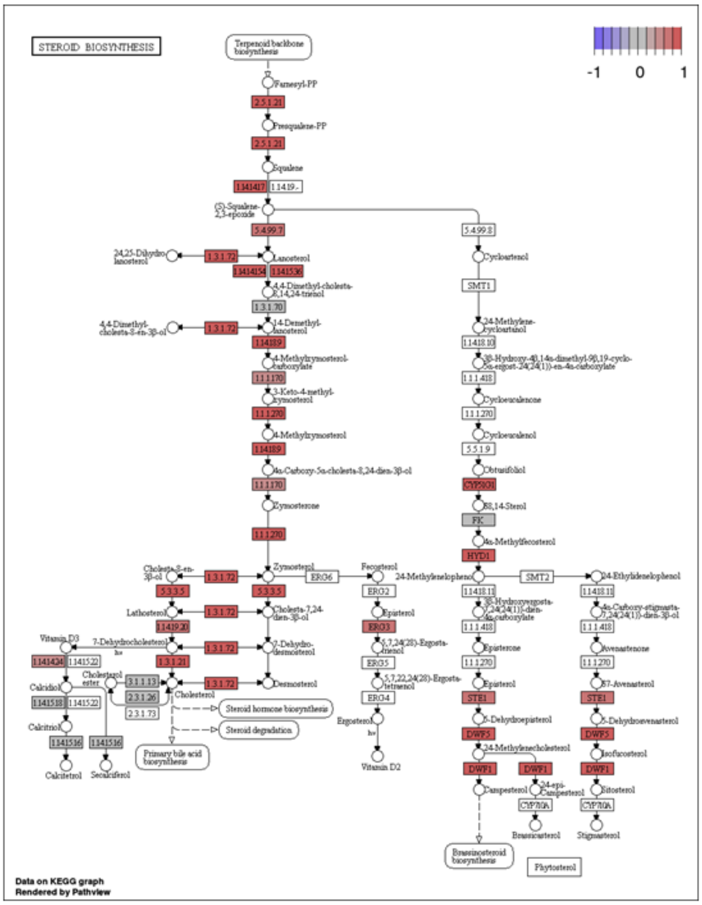


**B.** **CRISPR-ESC model**

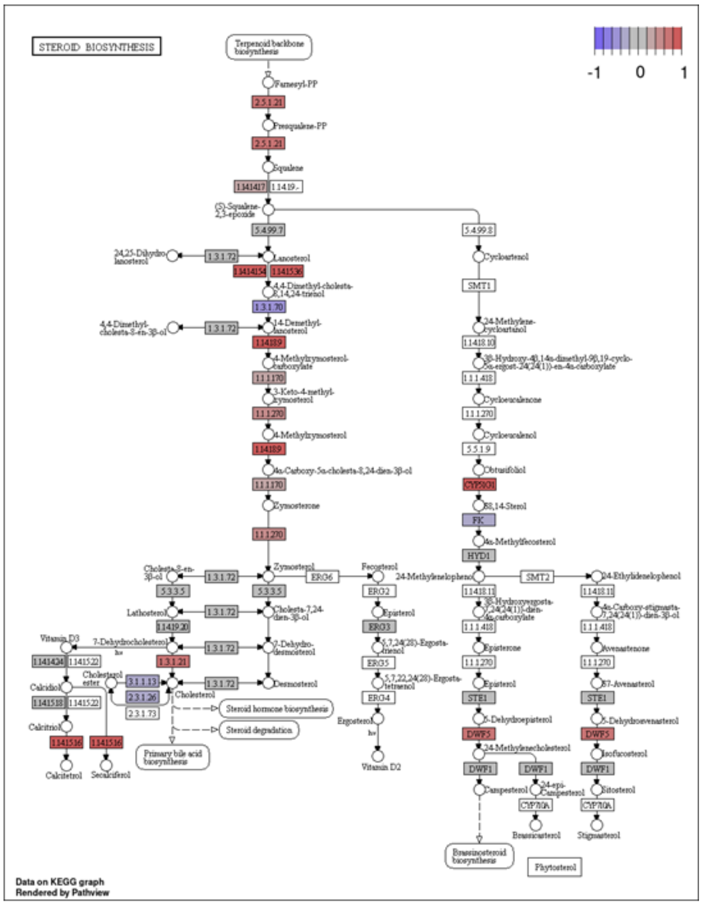


**C.** **Pretemer et al**

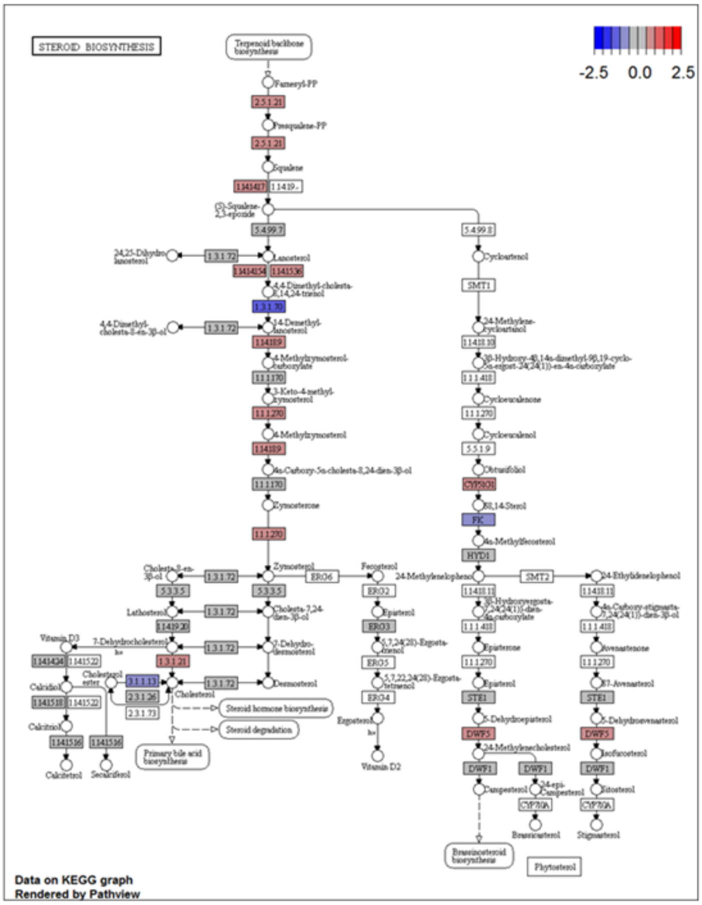


**D.** **Mouse model**

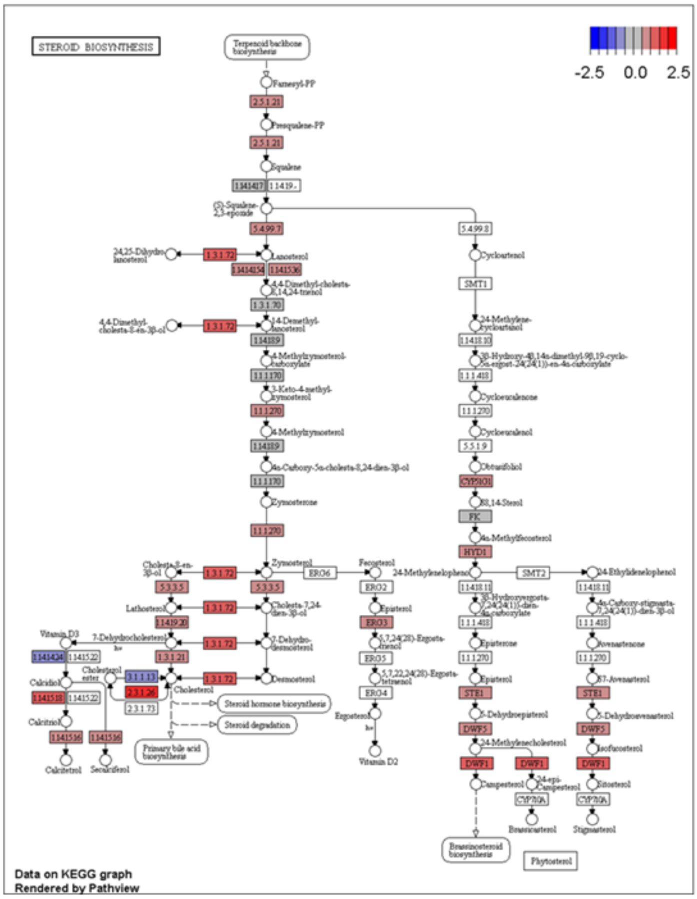

Figure S14. Pathview of cholesterol pathway in four models of MED. A, cholesterol pathway pathview in patient-iPSC model. B, cholesterol pathway pathview in CRISPR-ESC model. C, cholesterol pathway pathview in Pretemer et al model. D, cholesterol pathway pathview in mouse model. Colour indicated log2FC change in gene express for affected/mutant vs unaffected/WT for each of the four models. Red indicated increased gene expression, blue indicated decreased gene expression.
